# A coarse-grained model of glycosaminoglycans for biomolecular simulations

**DOI:** 10.1101/2023.10.06.561318

**Authors:** Aishwary T. Shivgan, Jan K. Marzinek, Alexander Krah, Paul Matsudaira, Chandra S. Verma, Peter J. Bond

## Abstract

Proteoglycans contain glycosaminoglycans (GAGs), negatively charged linear polymers made of repeating disaccharide units of uronic acid and hexosamine units. They play vital roles in numerous physiological and pathological processes, particularly governing cellular communication and attachment. Depending on their sulphonation state, acetylation, and glycosidic linkages, GAGs belong to different families. The high molecular weight, heterogeneity, and flexibility of GAGs hampers their characterization at atomic resolution, but this may be circumvented via coarse-grained (CG) approaches. In this work, we report a CG model for a library of common GAG types in their isolated or proteoglycan-linked states compatible with the widely popular CG Martini forcefields (versions 2.2 and 3.0). The model reproduces conformational and thermodynamic properties for a wide variety of GAGs, as well as matching structural and binding data for selected proteoglycan test systems. The parameters developed here may thus be employed to study a range of GAG-containing biomolecular systems, benefitting from the efficiency and broad applicability of the Martini framework.

## 1 Introduction

Proteoglycans (PGs) are heavily glycosylated proteins. These high molecular weight biomolecules found in the extracellular matrix (ECM) play vital roles in governing cellular interactions, the mechanical and viscous properties of tissues, and numerous physiological and pathological processes^1,2^. PGs bind to water, ions, chemokines, growth factors and many other biomolecules^2^. These properties of PGs arise from the covalently attached linear anionic glycosaminoglycans (GAGs), composed of repeating disaccharide units of uronic acid (UA) and hexosamine (HEX) units. The UA units can either be β-D-glucuronic acid (Glc) or α-L-iduronic acid (IdoA), while the hexosamine units can be α-D- or β-D-glucosamine (GlcN) or N-acetyl-β-D-galactosamine (GalNAc). Depending on their sulphonation state, acetylation, and glycosidic linkages, these building blocks make up various GAG families. These encompass non-sulphated GAGs such as hyaluronic acid (HA) and heparin (HP), and sulphated forms including heparin sulphate (HS), chondroitin sulphate (CS), dermatan sulphate (DS) and keratan sulphate (KS) (**Figure 1**). These GAGs are typically linked to a tetrasaccharide GlcA(β1→3)Gal(β1→3)Gal(β1→4)Xyl(β-Ser) sequence made of xylose (Xyl), two Galactose (Gal) units and Glucose (Glc), which then connects to the protein via an O-linked serine or threonine^3^.

**Figure 1.**
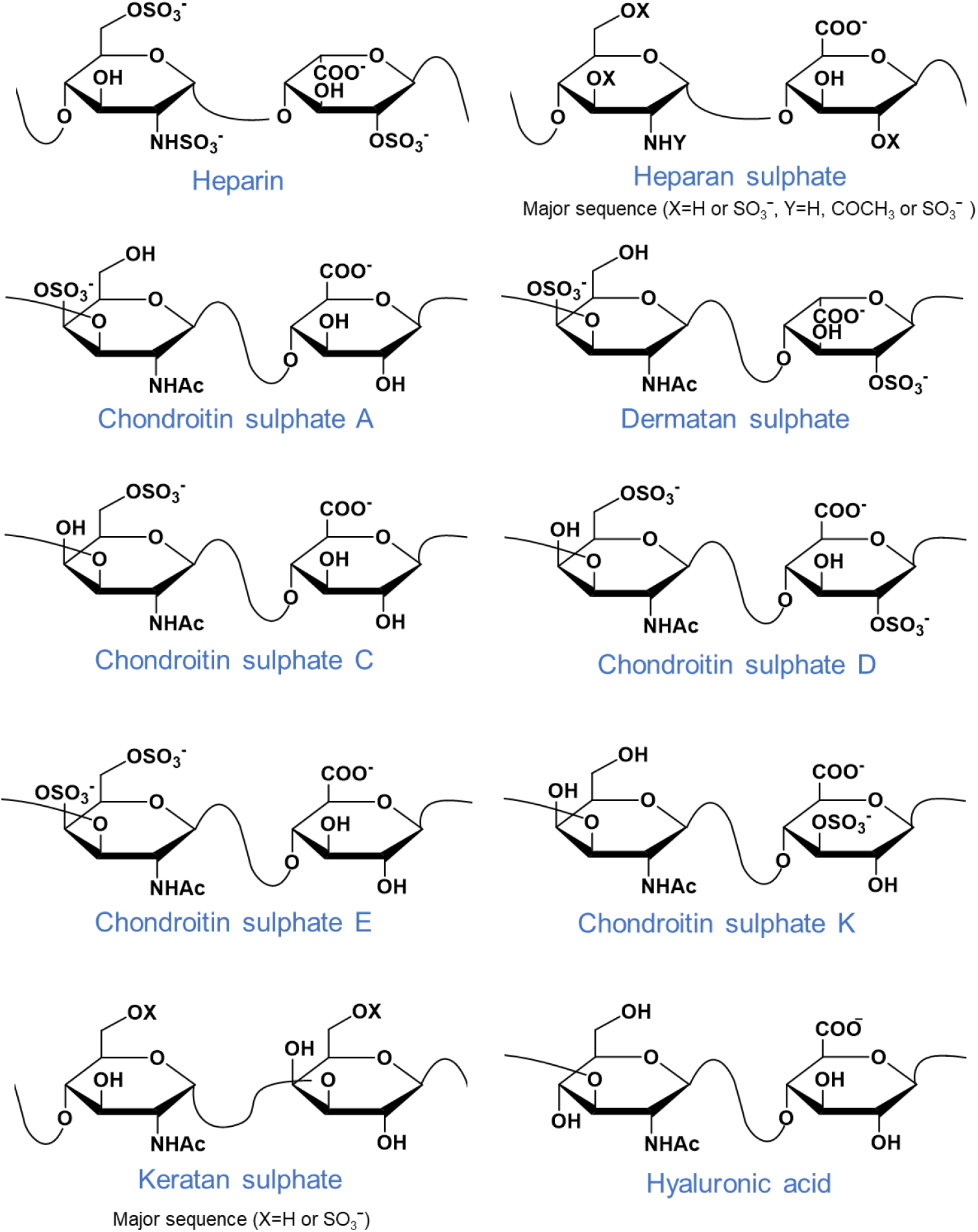
Classification of GAGs. The figure shows the major GAG structures for Heparin [IdoA2S(α1→4)**GlcNS6S(α1→4)IdoA2S(α1→4)**GlcNS6S], Heparan sulfate [GlcA(β1→4)**GlcNAc(α1→4)GlcA(β1→4)**GlcNAc], Chondroitin sulfate [GlcA(β1→3)**GalNAc4S(β1→4)GlcA(β1→3)**GalNAc4S], Dermatan sulfate [IdoA(α1→3)**GalNAc4S(β1→4)IdoA(α1→3)**GalNAc4S], Keratan sulfate [Gal6S(β1→4)**GlcNAc6S(β1→3)Gal6S(β1→4)**GlcNAc6S], Hyaluronic acid [GlcA(β1→3)**GlcNAc(β1→4)GlcA(β1→3)**GlcNAc]

GAGs are implicated in various disease states including inflammatory conditions^4^, cancers^5–7^, corneal opacification^8,9^, arthritis^10^ as well as age-related diseases^11^. Furthermore, GAGs in the glycocalyx surrounding cells can serve as attachment factors or receptors for infectious pathogens^12^. For example, host HS may stabilize the so-called ‘up’ conformation of the SARS-CoV-2 spike protein in concert with its own viral N-glycans, thereby facilitating angiotensin-converting enzyme 2 (ACE2) receptor binding and subsequent COVID-19 infection^13^. Unsurprisingly, there is significant therapeutic interest in GAGs, but experimentally studying their conformation, dynamics, and interactions at the molecular level is hampered by challenges associated with GAG synthesis^14^ and further complicated by their polymeric nature and the flexibility of their glycosidic linkages^15^. As an alternative, computational approaches may be utilized to better understand their structure-function relationships. The large size of polymeric GAGs makes quantum mechanics (QM) based methods computationally challenging, but their simulation is more tractable using molecular dynamics (MD) simulations. In MD, the nature and the accuracy of the atomic interactions are defined by bonded and non-bonded interaction potentials collectively termed the force field (FF). Such FFs are typically optimized based on QM and experimental data^14,16^. Many all-atom (AA) FFs such as GLYCAM^17^, CHARMM^16^ and GROMOS^18^ can correctly reproduce the conformational properties of GAG sequences and their protein binding properties^19,20^. AA studies have enabled e.g., improved understanding of anomeric effects or torsional preferences of sulphate groups, and characterization of critical inter- and intramolecular interactions that affect the flexibility and stability of GAGs^20,21^. Despite the high resolution accessible to AA FFs, their application to GAG sequences has typically been limited to, at best, dodecasaccharides, and in many cases, to tetra- or hexa-saccharides, due to the computational cost associated with obtaining sufficient sampling^22^. A solution to this is the development of simplified coarse-grained (CG) models, potentially enabling the study of conformational dynamics of higher-order GAG sequences as observed in nature (i.e. ∼20-200 disaccharides long).

In the CG approach, sets of atoms are grouped together and replaced by ‘pseudo-atomic’ particles, resulting in a reduced number of degrees of freedom in the system, thereby allowing access to longer simulation and time- and length-scales at the expense of molecular detail. One of the first CG models for CS and HA was developed by Bath et al., which could predict the influence of pH and ionic strength on persistence length and end-to-end distances, respectively^23^. The model was characterized by three beads representing a monosaccharide with an additional virtual particle representing its center of geometry and charge. In another CG model, Sattelle et al. implemented ring puckering in HP, CS and DS models to study their 3D shape, bioactivities and hydrodynamic properties^24^. In addition, the chain volume and its rigidity were shown to depend on glucuronic acid, the conformational properties of which could be modulated by the sulphonation state. An AMBER-compatible CG FF developed by Samsonov et al. used unique particle types based on several protein-GAG complexes^25^. In their model, the bonded interactions were optimized based on AA MD simulations, while the non-bonded parameters were optimized based on potential of mean force calculations. The resultant CG model reproduced many local and global properties from AA simulations, such as mean distributions of the root mean square deviation (RMSD), end-to-end distances, and radii of gyration (R_g_). In a CG model by Kolesnikov et al., a polymeric field-theoretical approach could reproduce osmotic pressure data and many solution properties in the presence of ions^26^.

Although CG models for GAGs alone exist, they are not necessarily compatible with other essential biomolecules such as proteins, lipids, RNA, or DNA, hampering the capacity to study GAGs in realistic, complex biological environments. Martini is currently the most widely used CG FF for biomolecular systems^27^. This FF has been designed to reproduce thermodynamic properties and partitioning data by using a few building blocks that mimic the chemical specificity of the underlying atoms^27^. In the Martini model, four heavy atoms are typically represented as a single CG bead (4:1 mapping). In addition, finer mappings, including 3:1 and 2:1, are utilized for representing rings or high-resolution features. A typical uncharged bead of CG Martini water represents four AA water molecules. A three-bead polarizable water model is also available, two of which are dipoles connected to the central bead^28^. Implicit solvation is also supported, called ‘Dry’ Martini, and currently is only compatible with lipid systems^29^. The modular building blocks, along with a small number of parameters, result in broad applicability and transferability to other classes of biomolecules such as proteins, lipids, DNA, and N-glycans^30–34^. Since its introduction, the model has seen wide adoption among the scientific community for studying problems like lipid self-organization, membrane fusion, protein oligomerization, drug delivery, and many more, as summarized elsewhere^35^. The combination of the Martini FF with advances in computational resources and multiscale approaches have made possible simulations of large and complex biomolecular systems such as mitochondria^36^, liposomes^37^ and virus particles such as dengue^38^ and influenza A^39^. Certain shortcomings in version 2.2 of the Martini FF have surfaced in recent years such as overly strong or ‘sticky’ interactions for proteins^40^ and carbohydrates^34^. In an attempt to alleviate these problems, Martini v3.0 was introduced, where rebalancing of the non-bonded interactions was implemented^41^, while new bead types and an expanded interaction matrix resulted in a wider chemical space.

In the present work, we have extended the Martini FFs (both v2.2 and v3.0) towards the simulation of various GAG molecules, including HP, HS, CS, DS, and HA, and their O-linked proteoglycan variants. The bonded parameters for GAGs, including linkers, were obtained from extensive AA simulation sampling. Multiple mapping schemes were tested for heparin and the most successful model was employed for other GAGs. Bead types for monosaccharides were validated by calculating partition coefficients and comparing them to predictions from various empirical methods. The resultant CG models were used to predict local and global GAG properties and were applied to some key case studies based on availability of relevant experimental data in the literature. Firstly, HP is essential for homo- and heterodimerization of the fibroblast growth factor receptor (FGFR), and GAG binding modes from simulations were compared with high-resolution structural studies and used to confirm its previously proposed capacity for binding longer HP ligands. Secondly, bikunin is a serine protease inhibitor used to treat acute pancreatitis, and its PG form contains a single O-linked CS chain that is essential for function; simulations of the PG complex were carried out to characterize the dynamics of the GAG moiety, validated via hydrodynamic size estimates. The CG GAG library developed here represents a computationally efficient strategy for delineating the biological roles of GAG molecules, GAG-protein complexes, and PGs.

## 2 Methods

### 2.1 Simulation parameters

In AA simulations, the GROMOS54a7_GLYC18_ united-atom FF was used for the UA and HEX units involved in various GAG monosaccharides and Xyl-β-Ser O-glycosidic linkages representing the tetrasaccharide GAG linker. The GAG molecules were solvated using the SPC water model^42^. The systems were neutralised using Na^+^ and Cl^-^ counterions. The simulations were performed at a temperature of 310 K using the velocity rescale thermostat^43^ with a relaxation time of 0.1 ps. The pressure was maintained at 1 bar using the weak coupling Berendsen barostat with a relaxation time of 1 ps. A cutoff of 1.4 nm was used for electrostatic and van der Waals interactions. Long-range electrostatics were treated using the particle mesh Ewald^44^ method with a cutoff of 1.2 nm.

In the case of CG simulations, both Martini v2.2^27,30,45^ and v3.0^41^ FFs were used. Systems were solvated using Martini water beads, and for systems using v2.2 FF an additional 10% antifreeze water beads were added. A temperature of 310 K and pressure of 1 bar were maintained using the velocity rescale thermostat^46^ and Parrinello-Rahman barostat^47^ with relaxation constants of 1 ps and 12 ps, respectively. The reaction-field method^48^ was used for electrostatics with a cutoff of 1.1 nm. Both AA and CG systems were minimized for 5,000 steps, equilibrated for 50 ns and then subjected to productions runs (summarised in **Table S1Error! Reference source not found.**).

GROMACS 2018^49^ was used for all the simulations and run on: i) an in-house Linux cluster using 20 CPUs (E5-2690v3) and 1 GPU (NVIDIA K40) and ii) the ASPIRE 1 supercomputer of the National Supercomputing Centre Singapore (https://www.nscc.sg) using 24 CPUs (E5-2690v3) and 1 GPU (NVIDIA K40).

### 2.2 Mapping schemes

Two mapping schemes were initially tested for the assessment of the stability of GAG chains (**Figure 2**). Linear (‘map1’) and ring-like (‘map2’) mapping schemes were tested for HP. The linear mapping scheme is similar to that previously developed for carbohydrates^30^ and N-glycans^34^. The ring-like scheme has previously been used for glycolipids^33^. The underlying chemical nature of the grouped atoms and previously modelled glycans were considered when selecting the bead types. In both schemes, charged functional groups (i.e. sulphates and carboxylates) were treated with a single negatively charged bead. Initial bonded parameters were obtained from tetrasaccharide simulations.

**Figure 2.**
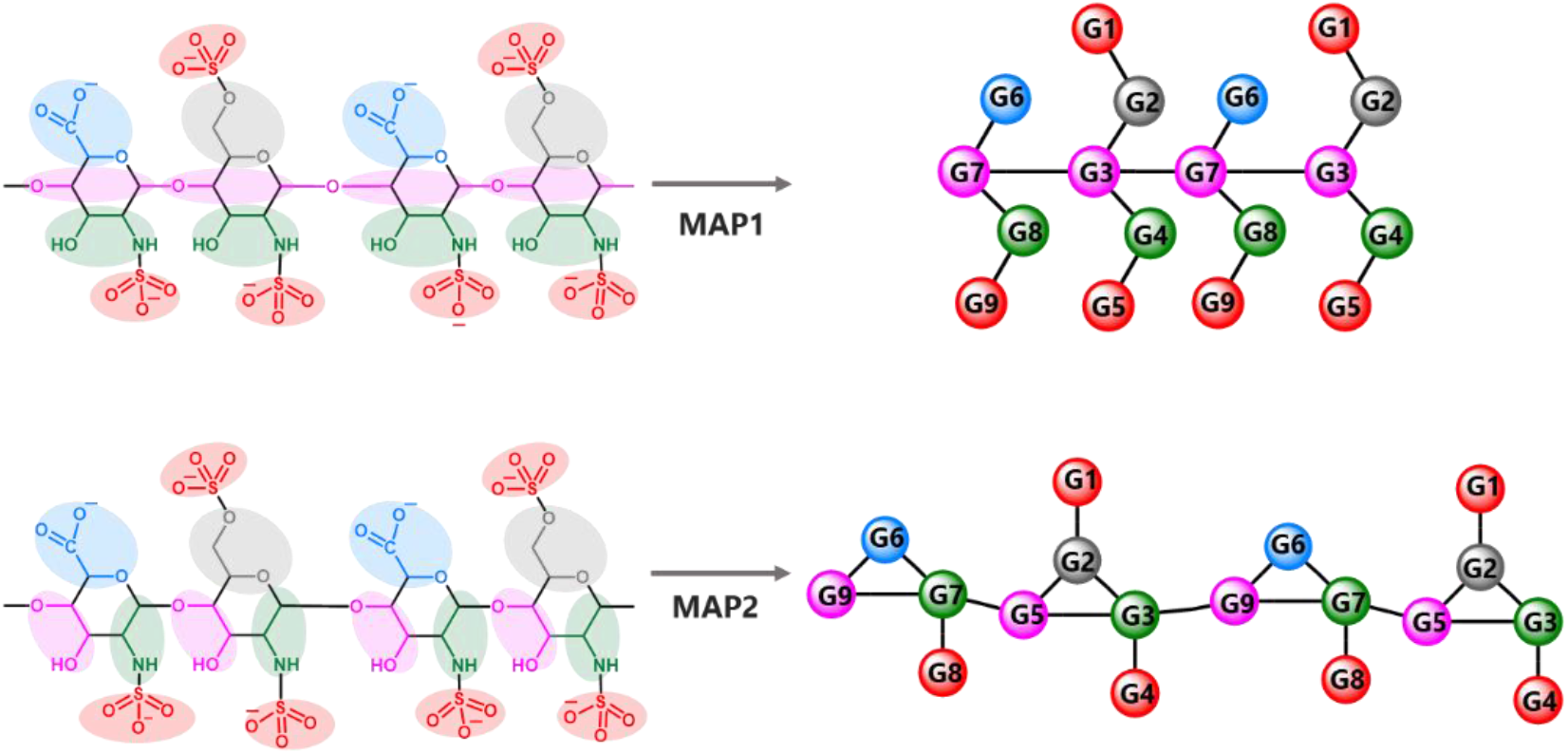
Mapping schemes for heparin. The first scheme makes a linear linkage for the monosaccharides, while the second scheme uses a ring-like topology represented by three beads. Sulphate and carboxylate groups are represented as separate beads in both schemes.

The bonds between beads were fitted using:

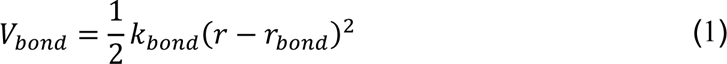

where *r_bond_* and *k_bond_* are the equilibrium distance and the force constant, respectively. The angles within the CG beads were treated using:

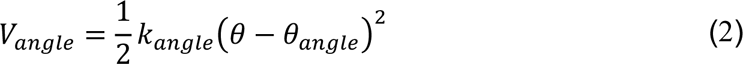

where *θ_0_* is the equilibrium angle and *k_angle_* is the force constant. The dihedrals were described using:

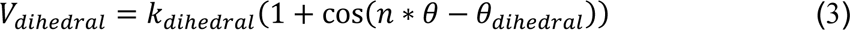

where *θ_dihedral_* is the equilibrium angle between planes defined by the coordinates of the four consecutively bound atoms *i, j, k* and *j, k, l* respectively, *k_dihedral_* is the force constant, and *n* is the multiplicity. The dihedral distributions had a single minimum and were fitted using a multiplicity of 1.

#### 2.2.1 Validation of non-bonded parameters

The initial bead types were selected by chemical intuition and by analogy with previously parametrized carbohydrate and N-glycan molecules^30,34^. The UA and HEX units had not been parametrized before. The additional sulphate groups make it necessary to validate the choice of the Martini bead types. As Martini parameterization is based on the reproduction of experimental partitioning coefficients (log *P*_*ow*_), these were calculated for various disaccharide combinations commonly found in GAGs. These coefficients were obtained using thermodynamic integration (TI) calculations with 55 windows along with additional windows around the high curvature region. Softcore potentials by Beutler (implemented in Gromacs) were used for sampling the LJ potential at high scaling values^50^. The calculations were performed in 1-octanol and water phases. The difference in solvation free energies of GAG disaccharides in 1-octanol (Δ*G*_*o*_) and water (Δ*G*_*w*_) was used to calculate the partition coefficients using ΔΔ*G*_*ow*_ = −2.3*RT* log *P*_*ow*_. The water phase was simulated using one GAG disaccharide solvated in a box of 1,000 water molecules, while the organic phase consisted of 43 water and 519 1-octanol molecules, representing a 0.255 water-octanol molar fraction^51^. The coefficients were obtained for both v2.2 and v3.0 versions of the Martini FFs. The obtained simulation data was compared to the results of empirical (log *P*_*ow*_) prediction methods within the ALOGPs 2.1^52^ program. The predicted coefficients from multiple methods such as ALOGPs, ALOGP, KOWWIN, ClogP, MLOGP, miLogP, AB/LogP. These methods differ from one another in terms of molecule fragmentation patterns and atomic contributions to the predicted log *P*_*ow*_. The choice of Martini beads for mapping the GAGs (based on the ‘map1’ scheme) is summarized in **Table 1**.

**Table 1.** Martini bead type selection for different CG GAGs as shown in Figure 4. The selections shown are for Martini v3.0. In the case of Martini v2.2, the Q1 bead was replaced with Qa bead. Mapping is based on ‘map1’ scheme (refer to **Figure 2**).

| GAG | Martini bead type |  |  |  |  |  |  |  |  |  |  |  |
| --- | --- | --- | --- | --- | --- | --- | --- | --- | --- | --- | --- | --- |
|  | G1 | G2 | G3 | G4 | G5 | G6 | G7 | G8 | G9 | G10 | G11 | G12 |
| HP | Q1 | P1 | SP2 | P2 | Q1 | Q1 | SP2 | P2 | Q1 |  |  |  |
| HS | Q1 | P1 | SP2 | P2 | Q1 | Q1 | Q1 | SP2 | P2 | Q1 |  |  |
| CS | Q1 | P1 | SP2 | P2 | SP1 | Q1 | Q1 | SP2 | P2 | Q1 |  |  |
| DS | P1 | SP2 | P2 | SP2 | Q1 | Q1 | SP2 | P2 | Q1 |  |  |  |
| KS | Q1 | P1 | SP2 | P4 | SP2 | Q1 | P1 | SP2 | P4 |  |  |  |
| HA | P1 | SP2 | P2 | SP1 | Q1 | SP2 | P4 |  |  |  |  |  |
| LIN | SP1 | SP2 | P4 | P1 | SP2 | P4 | P1 | SP2 | P4 | Q1 | SP2 | P4 |

#### 2.2.2 Protein-GAG studies

The AA crystal structures of FGFR1-FGF2^53^ dimer (PDB:1FQ9) and bikunin^54^ (PDB:1BIK) were converted to CG representation using the *martinize.py* script available at cgmartini.nl. An elastic network was applied for preserving higher-order protein structure. The network was applied to atoms within the lower and upper cutoff distances of 0.5 and 0.9 nm with a force constant of 500 kJ mol^-1^ nm^-2^. These values ensured that the folds of each protein complex were retained, with the backbone RMSD of each system with respect to the experimental structure plateauing at ∼5 Å by the end of each trajectory (**Figure S1)**.

In the case of a HP20 GAG pre-bound to the FGFR1-FGF2 complex, the GAG was manually aligned to the GAG HP fragment co-crystallized with the X-ray structure (PDB:1FQ9) and simulated for 500 ns using the Martini v2.2 FF with the protein-to-GAG non-bonded LJ interactions scaled by 0.9. This scaling of LJ interactions has previously been shown to be important for correctly representing the solution behaviour of glycans and their interactions with proteins^34^. Besides this, multiple unbiased self-assembly simulations were run, based on two sets of initial configurations of a simulation box containing FGFR1-FGF2 and two randomly placed HP_20_ molecules. Two replicates of each of the configurations were simulated for 1 μs which resulted in 4 independent simulations. In the case of bikunin, the GAG is attached via an O-linked modification at the Ser-10 position^55^. The protein structure of bikunin (PDB:1BIK) has been well studied^54^ while the heterogeneous GAG sequences^56^ attached to the protein have been characterized and found to range from a length of 27-39 saccharide units^57^. Therefore, either a 30-mer CS (CS_30_) or a 40-mer CS (CS_40_) chain was attached to the O-glycosylation site at Ser10^58^. Both systems were solvated in a cubic box and production runs were performed for 5 μs.

## Results

### 2.3 Mapping scheme

Initially, AA tetrasaccharide simulations were used to seed bonded parameter estimates for simulations of longer GAG chains. In the case of HP_10_, the ‘map1’ scheme (**Figure 2**) resulted in a very stiff chain with higher end-to-end distances compared to AA data, and reducing the associated dihedral force constants did not lead to significant improvement. On the other hand, the ‘map2’ scheme (**Figure 2**) resulted in collapse of the polymer into an aggregated state, and consequently, very low end-to-end distances compared to AA data. In the latter case, only extremely strong (10-fold increase) dihedral force constants improved the end-to-end distances (**Figure 3**). Overall, the linear mapping scheme or ‘map1’ resulted in better correlations with the AA data in these preliminary tests.

**Figure 3.**
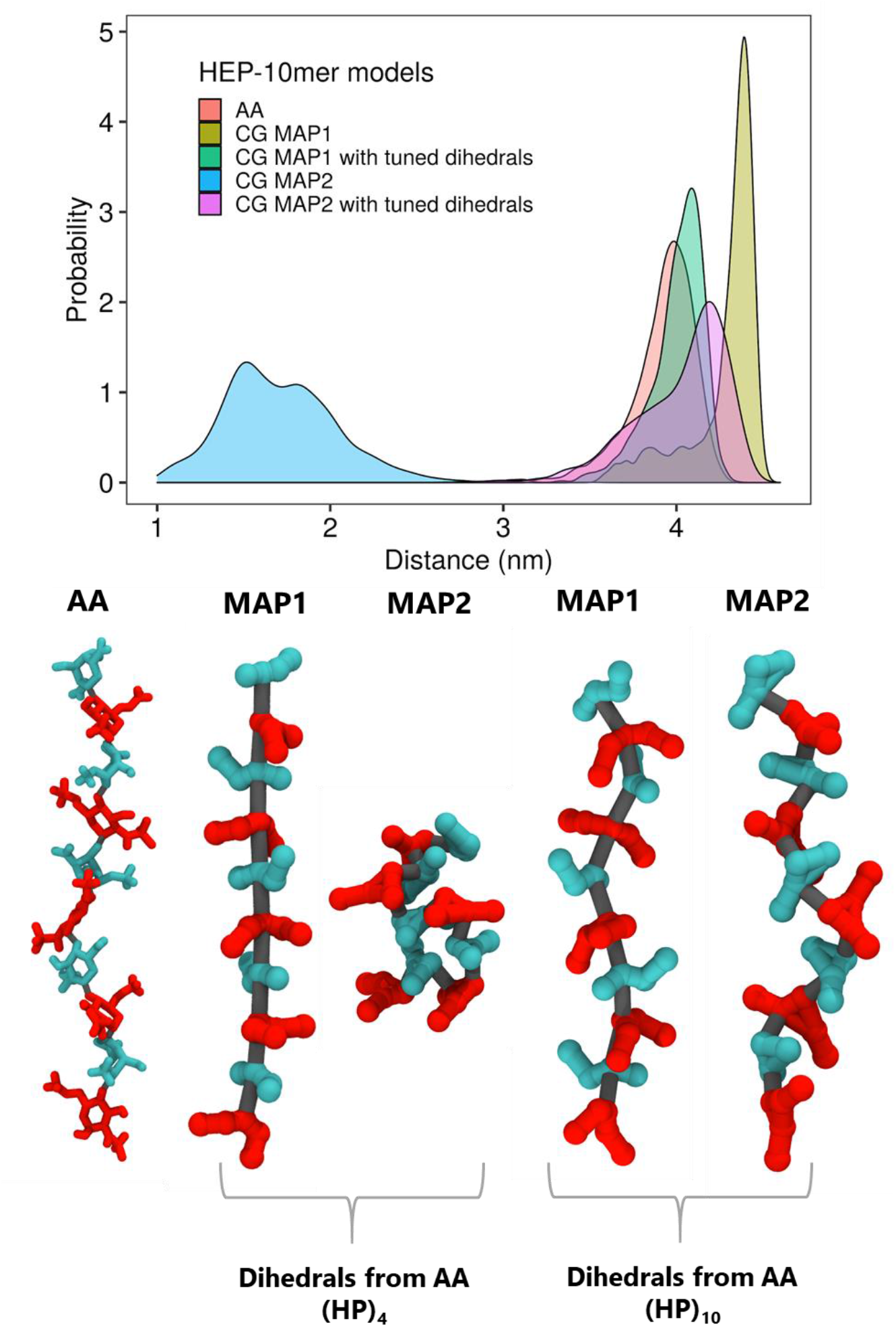
End to end distances of HP_10_ using two mapping schemes. (Top) Distribution of end to end distances. Distances were calculated from beads representing C1 and C4 atoms of the first and last monosaccharides. The AA trajectory was converted to a pseudo-CG trajectory to compare the end-to-end distances from the CG simulations. (Bottom) Final simulation snapshots of HP_10_ models. The hexosamine units are represented in red, while uronic acid units are colored in cyan.

To further improve the dynamics of longer GAG chains, backbone angles and dihedrals were next optimized against AA simulations of longer decasaccharide chains, initially focussing on HP_10_ (**Figure 3**). The backbone dihedral angles ϕ and ψ for beads connecting UA-HEX-UA-HEX and HEX-UA-HEX-UA sugars were monitored every GAG chain (**Figure 4** and **Figure 5**). For example, in the case of HP, these angles were defined for G7-G3-G7-G3 and G3-G7-G3-G7 beads (**Figure 4**). This led to an improvement in end-to-end distances and dihedral conformations, and reproduction of the helical nature of the chains (**Movie S1, Figure 5 and Figure 6**), resulting in a much better overall correlation between AA and CG data (**Figure 3**). The ‘map1’ scheme (**Figure 2**) was eventually selected as the model of choice and as the basis for subsequent parameterization of all other GAG molecules. This was because the ‘map2’ scheme required four backbone dihedrals to represent each tetrasaccharide unit, compared to two for ‘map1’, and with higher force constants that often resulted in simulation instabilities arising from constraint errors. Furthermore, visual inspection of simulated chains using the ‘map2’ scheme revealed unwanted monosaccharide rotations. The final set of refined bonded parameters are detailed in **Table S2**.

**Figure 4.**
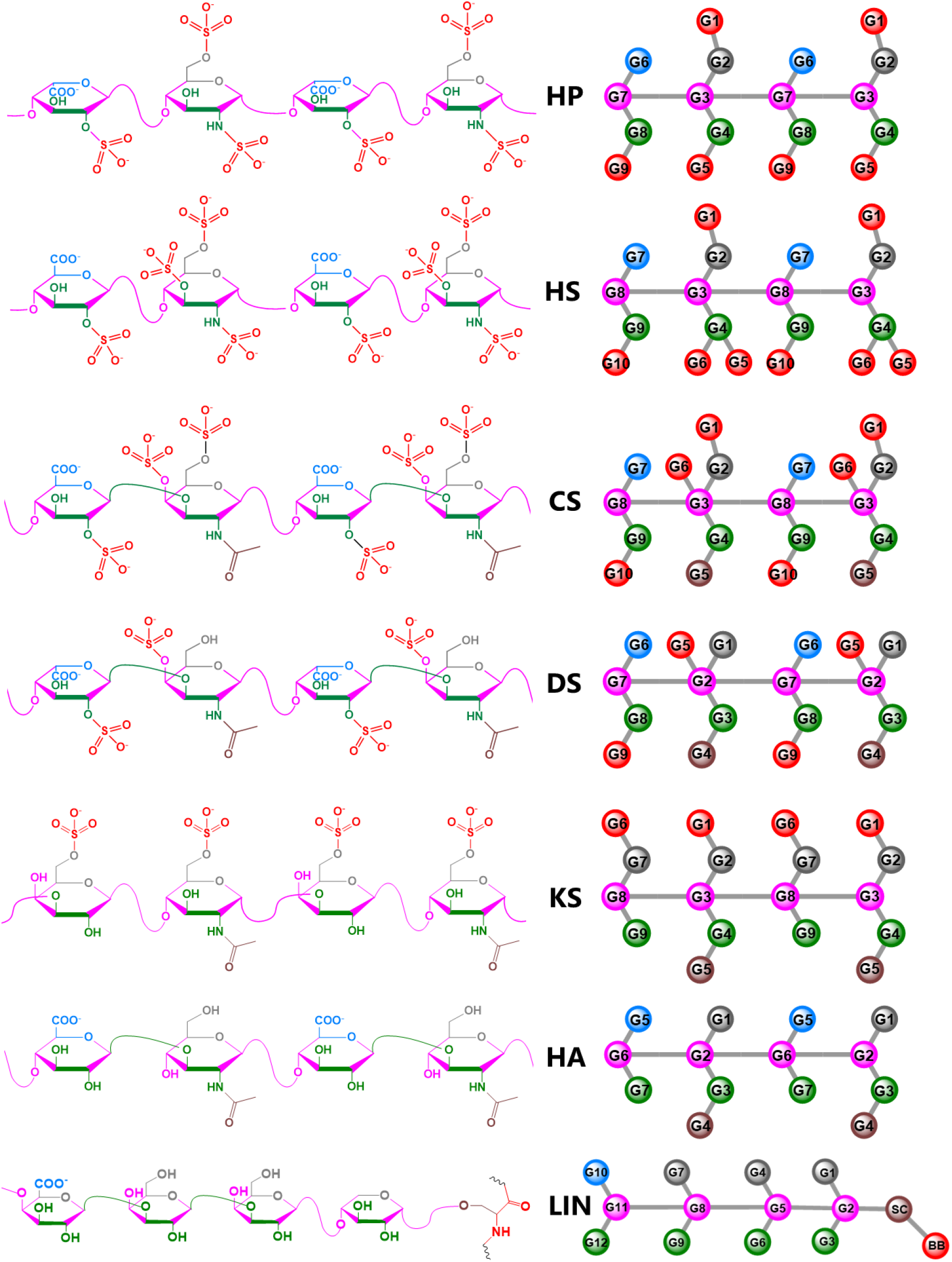
Mapping schemes for all GAG sequences. In addition to the common GAG sequences, the tetrasaccharide linker or **LIN** [GlcA(β1→3)Gal(β1→3)Gal(β1→4)Xyl(β-SER)] for the linkage between GAG and SER was parametrised.

**Figure 5.**
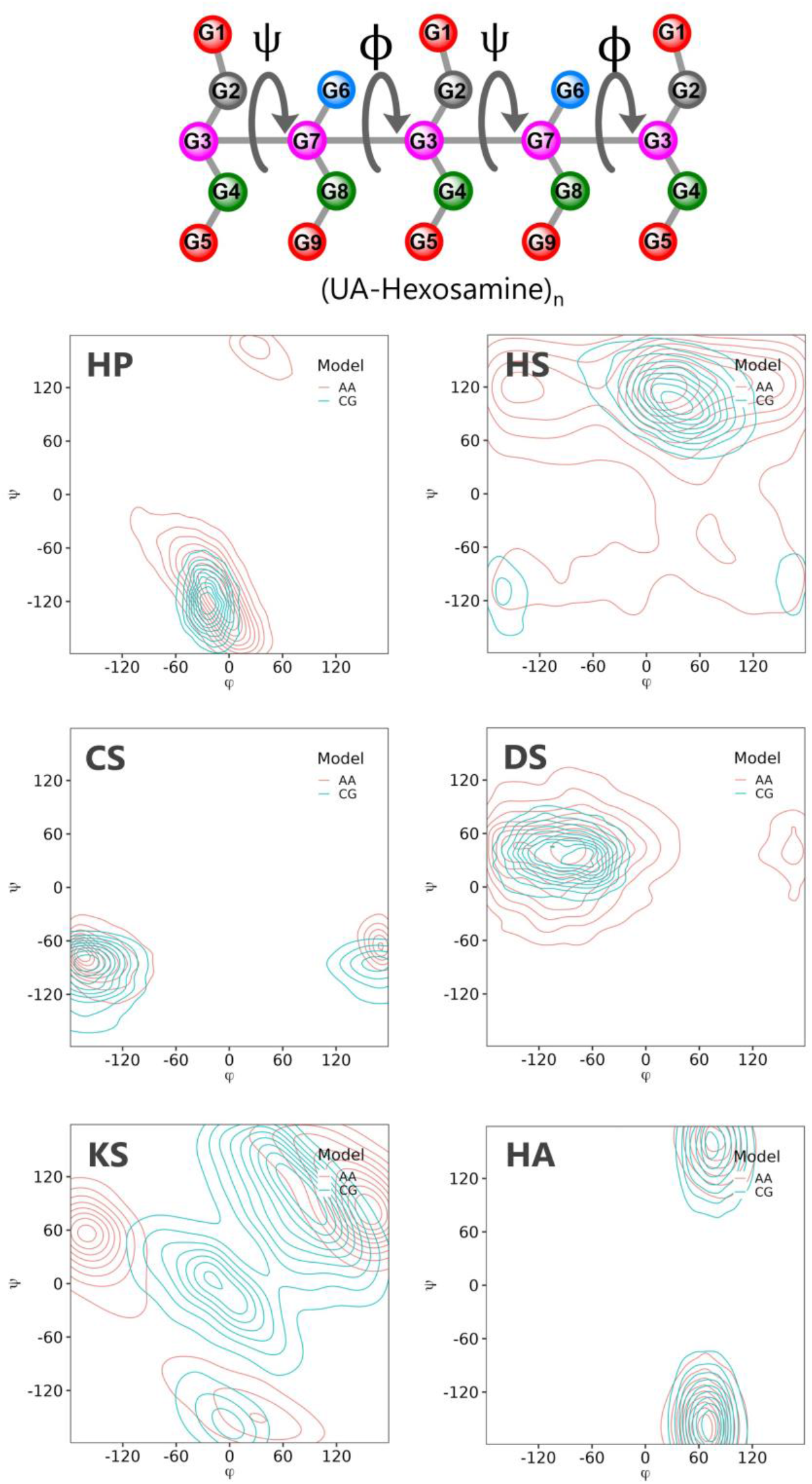
Backbone dihedral distributions of GAGs in atomistic and CG models. CG data was based on Martini v2.2. ϕ and Ψ angles were defined as dihedrals between neighboring disaccharide units in the CG model. For example, in the case of heparin, ϕ is defined as the dihedral angle between the G3-G7-G3-G7 beads while Ψ is defined as the angle between the G7-G3-G7-G3 beads. The angles were plotted as contour plots for HP, HS, CS, DS, KS and HA. The Martini bead definition is shown in **Table 2**.

**Table 2.** Hydrodynamic radii (*R_h_*) of GAG polymers from CG simulation and experiments.

| GAG | $R_h$ (sim) / nm | $R_h$ (DLS) / nm |
| --- | --- | --- |
| CS<sub>100</sub> | $4.8 \pm 0.3$ | $\sim 4.5^{61}$ |
| DS<sub>80</sub> | $3.8 \pm 0.2$ | $\sim 4.2^{61}$ |
| HP<sub>20</sub> | $1.9 \pm 0.03$ | $1.9^{60}$ |

**Figure 6.**
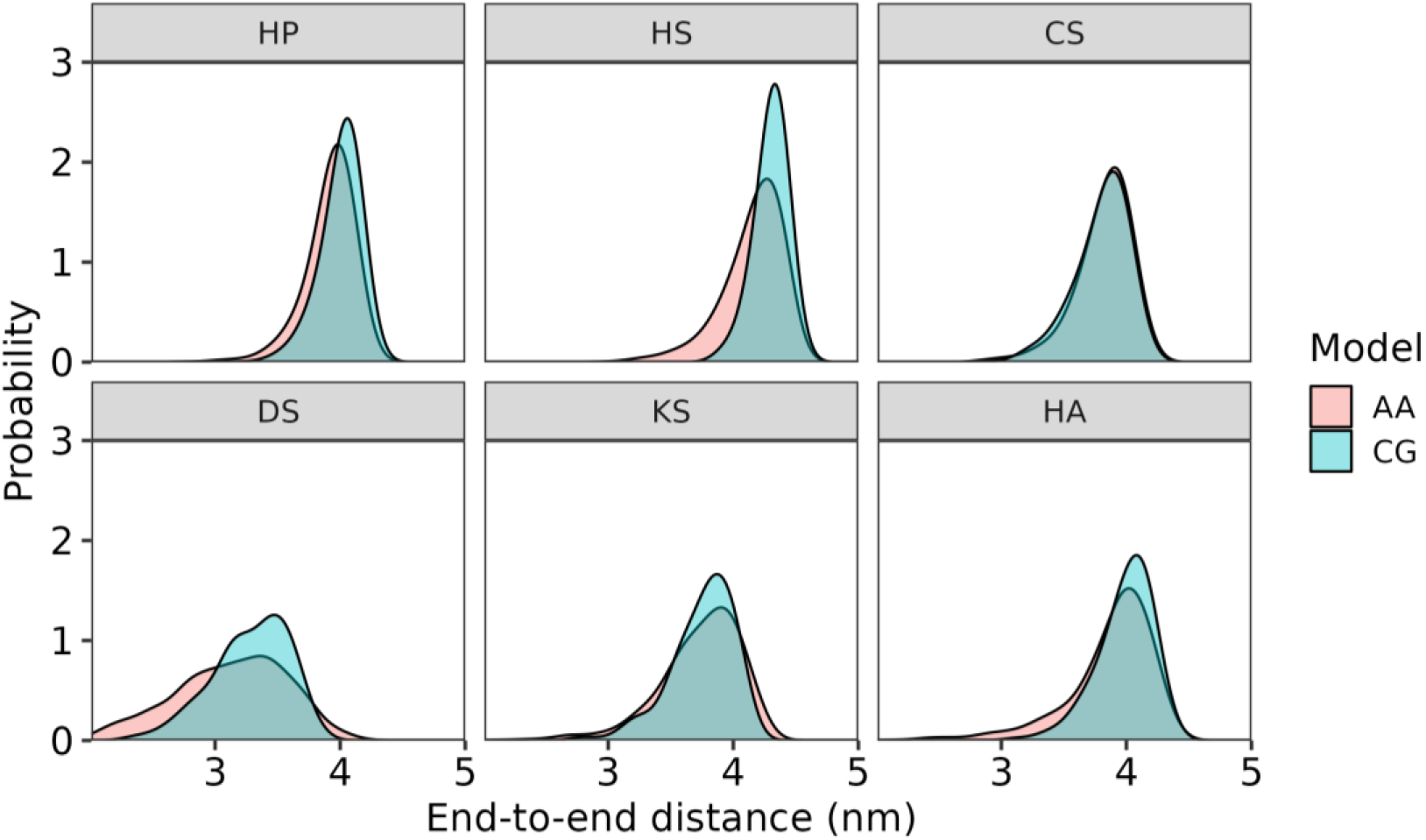
End to end distance distributions for decasaccharide GAG sequences. The distances were calculated from beads representing C1 and C4 atoms of the first and last monosaccharides. AA trajectories were converted to the pseudo-CG trajectories to compare with Martini v2.2 CG simulations.

### 2.4 Conformational properties of GAGs

The local conformational properties of CG 10mer GAG sequences using the selected linear ‘map1’ scheme were generally in good agreement with the AA data (**Figure 5**), as were end-to-end distances (**Figure 6**). In the case of CS and HA, the dihedral distributions were almost identical to their AA counterparts. The CG models of HP, HS and DS resulted in slightly narrower dihedral distributions, while correctly reproducing the AA equilibrium values. As shown for HP (**Figure S2**), lowering the dihedral force constants of the CG model resulted in deviation from the global AA minimum. In the case of KS, the AA distributions exhibited multiple local minima, which is problematic to achieve at CG resolution. Nevertheless, our final CG model enabled two of the three minima to be well sampled during simulations, while accurately reproducing the chain flexibility and end-to-end distances between the central bead of the first and last monosaccharide in KS_10_ (**Figure 6**).

### 2.5 Hydrodynamic radii

The hydrodynamic radius (*R*_ℎ_) describes the sphere of diffusion volume, helping to characterize polymer dynamics, and is typically similar to the radius of gyration (*R*_*g*_)^59^. We compared *R*_*g*_ values calculated from CG simulations of S_100_, DS_80_ and HP_20_ to available *R*_ℎ_values obtained from dynamic light scattering (DLS) experiments^60,61^. The *R*_*g*_was measured for the last 500 ns of each 1 μs simulation and used to estimate *R*_ℎ_ based on the relation *R*_*g*_ = 1.17 ∗ *R*_ℎ_ derived for polymeric chains^62^. These computed *R*_ℎ_ estimates for both the shorter and longer GAGs were in good agreement with experimental values (**Table 2**), consistent with the prior calibration of each GAG polymer in terms of both local structural/dynamic properties and end-to-end distances of up to 10mer sequences. The slight deviations observed for the longer CS_100_ and DS_80_ molecules may have been a result of simulating the fully sulphated forms, compared to the GAGs derived from natural sources used in experiments, which typically have variable sulphonation states that may affect chain flexibility and dynamics.

### 2.6 Validation of non-bonded parameters

The newly defined UA and HEX monosaccharides used across the various GAG sequences were validated by calculating partition coefficients (P_Oct→Wat_) for both Martini v2.2 and v3.0, based on a series of TI calculations as described in the Methods section. Solvation free energies for each GAG obtained in organic (Δ*G*_*oct*_) and water (Δ*G*_*wat*_) phases (**Table 3**) were used to obtain log *P*_*oct*_*_→_*_*wat*_ coefficients (**Table S3**). As expected, differences in solvation free energies were observed between the two Martini FFs. All values obtained for both Δ*G*_*oct*_and Δ*G*_*wat*_ were negative but were consistently greater in magnitude for v2.2 compared to v3.0 of the Martini FF, in agreement with the former being more ‘sticky’^41,63^. The negative ΔΔ*G*_*oct*_*_→_*_*wat*_ values obtained for both FFs reflect the preference for GAGs to partition into the water phase, due to the presence of polar hydroxyls along with anionic carboxylate and sulfate groups. Martini FF v3.0 exhibited slightly larger ΔΔ*G*_*oct*_*_→_*_*wat*_ values, indicating a greater tendency to partition into the bulk water phase.

**Table 3.** Solvation free energies calculated for GAG disaccharides with Martini FFs v2.2 and v3.0.

| GAG | Martini FF | $\Delta G$<br>(*Water*)<br>kJ mol<sup>-1</sup> | $\Delta G$ (*Octanol*)<br>kJ mol<sup>-1</sup> | $\Delta\Delta G$<br>(*Oct*→*Wat*)<br>kJ mol<sup>-1</sup> |
| --- | --- | --- | --- | --- |
| Heparin (HP) | v3.0 | -183.0 | -132.4 | -50.7 |
|  | v2.2 | -189.3 | -148.1 | -41.2 |
| Heparan-Sulphate (HS) | v3.0 | -200.6 | -146.7 | -53.9 |
|  | v2.2 | -202.0 | -157.3 | -44.7 |
| Chondroitin-Sulphate (CS) | v3.0 | -183.0 | -138.2 | -44.9 |
|  | v2.2 | -192.7 | -169.1 | -23.5 |
| Dermatan-Sulphate (DS) | v3.0 | -163.0 | -123.6 | -39.4 |
|  | v2.2 | -173.7 | -150.1 | -23.6 |
| Keratan-Sulphate (KS) | v3.0 | -164.7 | -121.5 | -43.1 |
|  | v2.2 | -186.1 | -150.6 | -35.5 |
| Hyaluronan (HA) | v3.0 | -120.94 | -91.35 | -29.6 |
|  | v2.2 | -144.55 | -118.76 | -25.8 |

As experimental values are unavailable, likely due to the highly polar nature of GAGs, the partition coefficients calculated from solvation free energies were compared to estimates obtained from empirical prediction methods including atomic (AlogP), hybrid (XlogP3) and fragment-based (ClogP, AC_logP) approaches (**Figure 7 and Table S3Error! Reference source not found.**). The predicted values correlated slightly better in Martini v3.0 than v2.2, wherein the average unsigned errors (AUE) compared to the mean of the empirical predictions were 8.14 and 8.5 kJ mol^-1^, respectively. The average of the predicted values from all these methods was generally slightly closer to the ones calculated from Martini v3.0 than to v2.2.

**Figure 7.**
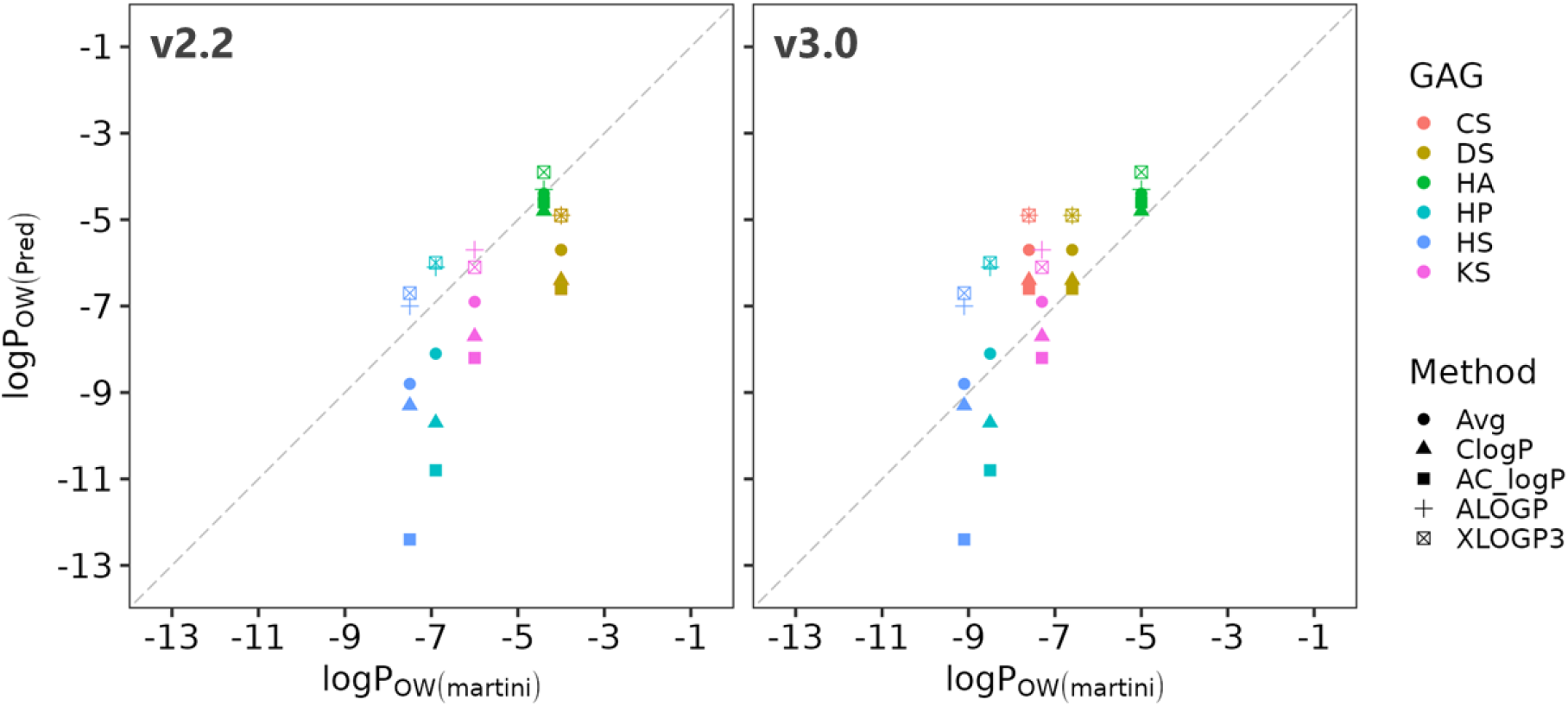
Partition coefficients (log *P_ow_*) of different GAG sequences. The log P_OW_ values obtained from simulations were compared to those obtained from empirical prediction methods as labelled. Avg represents the mean of values from all the prediction methods.

### 2.7 Aggregation studies

The highly polar and often charged nature of GAGs make them readily soluble in water. HP is water soluble with concentrations as high as 50 g L^-1^ ^64^. Therefore, we performed aggregation studies of 35 HP tetrasaccharides corresponding to a significantly lower concentration of ∼23 g L^-1^ (**Figure 8**). In Martini v2.2 simulations, the GAGs aggregated within the first few hundred nanoseconds and remained bound to one another upon extending the simulations to 1 μs. This suggested that the non-bonded interaction potentials are too attractive. Similar behaviour has previously been shown for carbohydrates^65^ and N-glycans^34^ when using the Martini v2.2 FF. To correct for the imbalance in the glycan-glycan nonbonded interactions, the well depths (ε) of the LJ interaction potentials were scaled according to the following equation:

**Figure 8.**
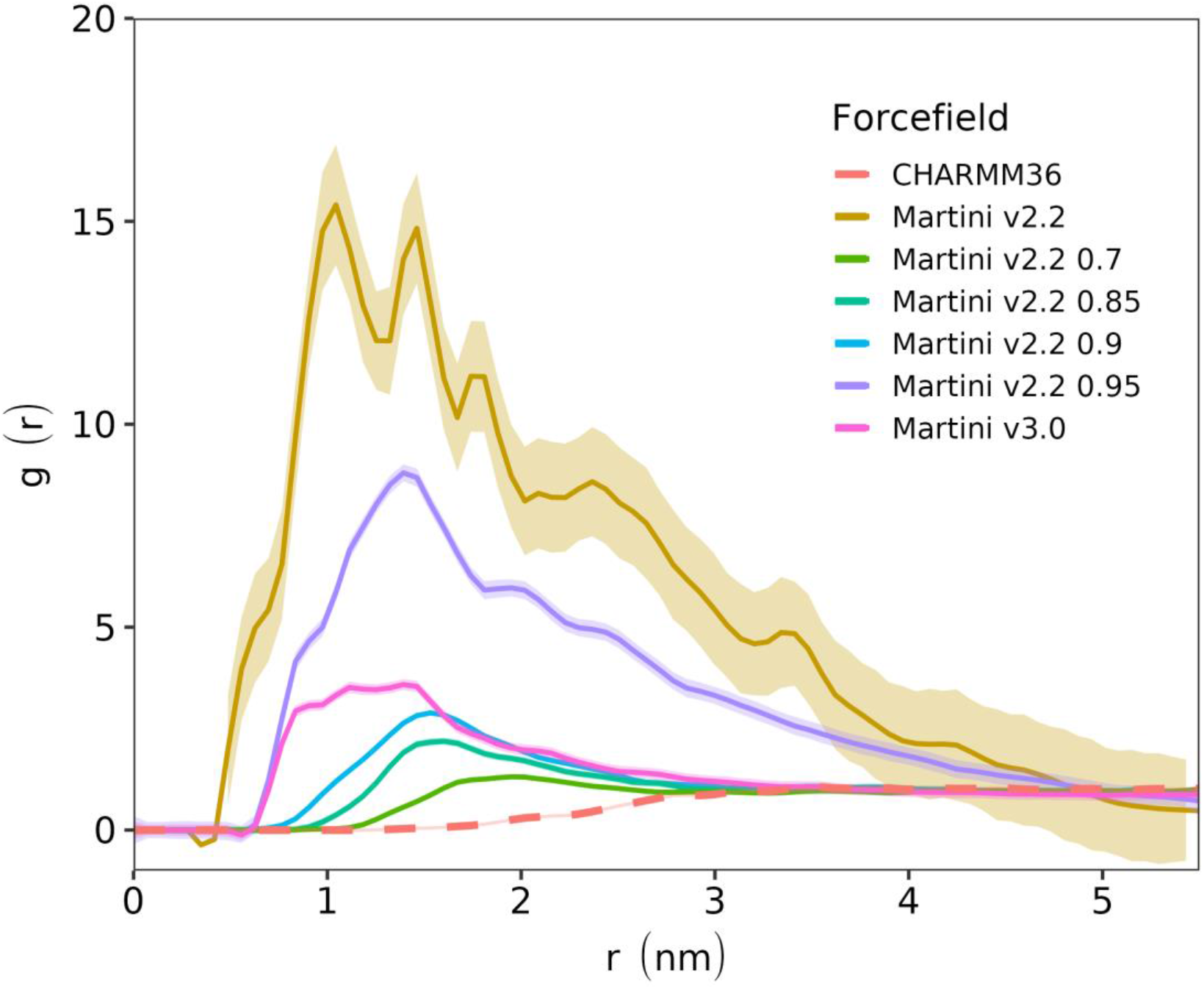
Aggregation propensity of heparin tetrasaccharides for different FFs and LJ scaling factors. The average RDFs for CG and AA models are shown. A range of FFs and LJ scaling factors were tested, as labeled. The RDFs were calculated from the final 200 ns of each trajectory.

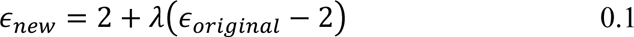

where *λ* is the scaling factor and *ε_original_* is the unscaled well depth of the LJ potential. The same correction factor has been used before for glycan^34,65^ and protein^40^ parameters. Multiple scaling factors, including 0.95, 0.9, 0.85 and 0.8 were tested. The radial distribution factions *g*(*r*) from CG simulations were compared to those obtained from AA simulations using the CHARMM36 FF (**Figure 8**). A drastic contrast in the *g*(*r*) values for unscaled Martini v2.2 simulations (*λ*=1.0) and the AA simulations was observed, suggesting excessive aggregation propensity. Scaling the interactions by *λ*=0.95 resulted in about 50% lower aggregation. Further scaling the interactions reduced the RDFs even more, and no aggregates were observed upon visual inspection of the trajectory in the case of *λ*=0.9. Although CG simulations could not perfectly reproduce the AA curve using *λ*=0.9 scaling or below, the differences between CG and AA models were minimal. Therefore, when using Martini v2.2, a scaling factor of 0.9 is recommended for GAG studies, similar to the value recommended for N-glycans^34^, where smaller glycans (up to 5 monosaccharides) needed the non-bonded interactions to be scaled by 0.9. In contrast, Martini v3.0 showed much lower aggregation propensity, as expected: the resultant RDF was similar to that obtained for Martini v2.2 with a scaling factor of 0.9.

### 2.8 GAG-protein studies

#### 2.8.1 Heparin-binding to FGFR1-FGF2 complex

FGFR1 is a dimeric receptor that is tightly regulated by a unique set of FGFs ligands^66,67^, and comprises immunoglobulin-like domains in the extracellular region, a transmembrane helix, and cytoplasmic domains, including a kinase domain^68,69^. HP is essential for the homo- and heterodimerization of the receptor, and hence crucial in the proper functioning of the signalling process^70,71^. GAG binding experiments with FGFR showed dependence on HP chain length. Hexasaccharides and even disaccharides exhibit some binding activity against the receptor^72–74^, while progressively increasing affinities have been reported for up to dodecasaccharides ^16,75^. Crystallographic studies showed that FGFR GAG binding is facilitated by a positively charged ‘canyon’ which may be occupied by two HP_10_ chains^53^ (**Figure S3Error! Reference source not found.**) with docking studies indicating possible occupation of the entire canyon by longer chains^71^. Therefore, we studied interactions of HP_20_ binding to the FGFR1-FGF2 complex via both biased docking approaches and unbiased self-assembly simulations (see Methods section for further details). In the pre-docked simulations, HP_20_ remained stably bound throughout the simulations (**Figure 9**). The saccharide units within the central cavity of D-II showed minimum fluctuations, as assessed visually, with terminal sugars showing higher flexibility (**Figure 9B**). The homodimeric symmetric interaction pattern of HP_20_ within the positively charged canyon was evident from contact probabilities averaged over the entire trajectory (**Figure 9C**), which was similar to that observed in the crystal structure.

**Figure 9.**
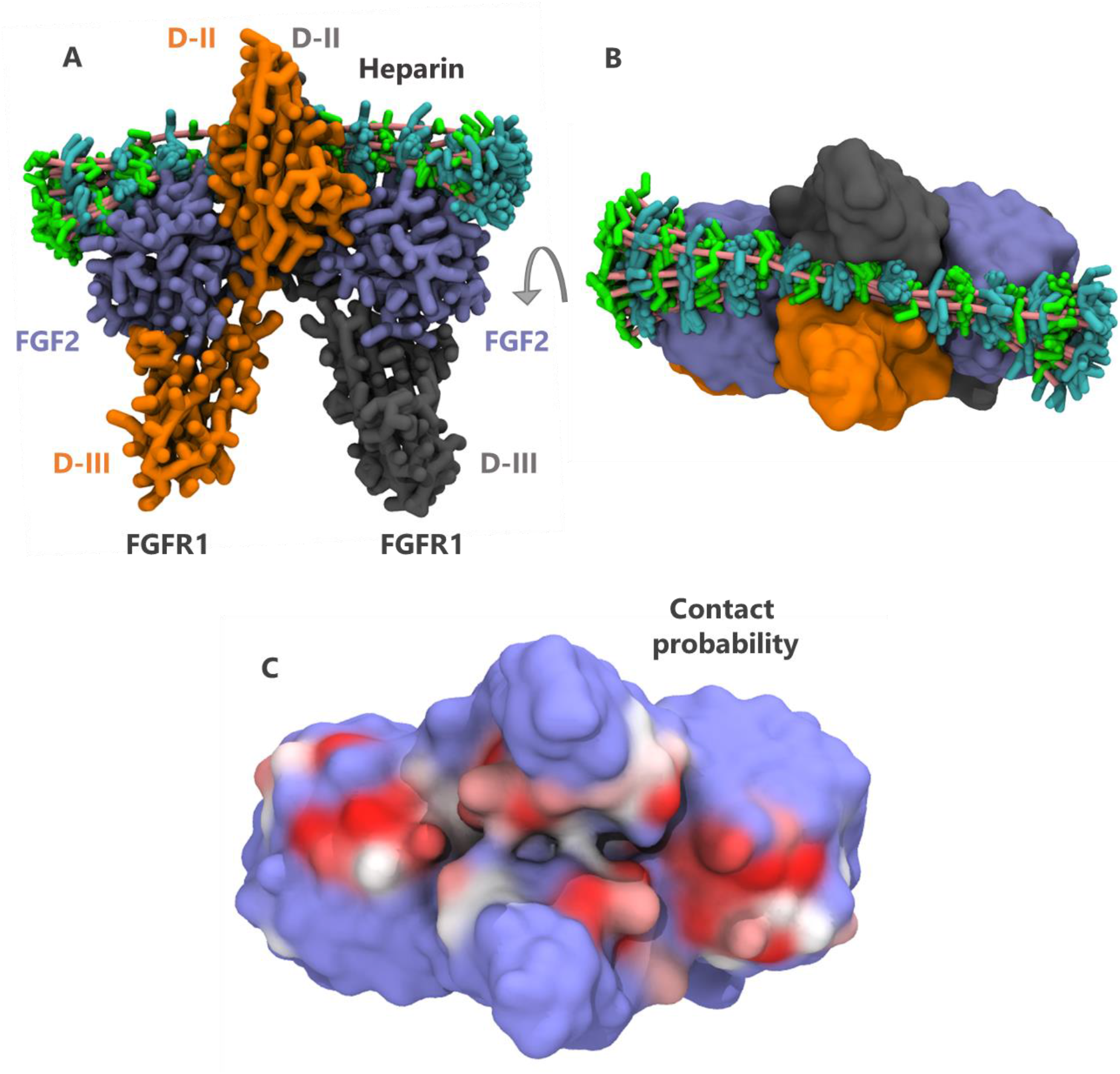
CG simulations of FGFR1-FGF2 complex with docked HP_20_. An ensemble of conformations of HP_20_ is shown as 20 overlapping and equally spaced frames derived from a 500 ns trajectory, viewed from **(A)** the side, and **(B)** the top of the complex. The GlcNS and IdoA sugars of HP_20_ are shown in licorice representation with cyan and green colors, respectively, and its backbone is colored salmon pink. Protein is colored as in Figure 8A-B. **(C)** Contact probability of HP_20_ to FGFR1-FGF2 complex (shown in surface representation) within a cutoff of 6 Å, colored on a scale from blue (P=0) through grey to red (P=1). Simulations were performed using Martini v2.2 parameters with a LJ scaling factor of 0.9.

As the CG representation allows access to higher timescales, unbiased self-assembly simulations of the receptor in the presence of two randomly placed HP_20_ ligands were next performed. In 3 out of 4 self-assembly simulations, HP_20_ ligands were able to find the binding site on FGFR1 within a few hundred nanoseconds (**Figures S4A-C**), with some HP_20_ saccharides binding to positively charged residues on FGF2 in the remaining replicate (**Figure S4D**). In one of the productive replicates, an HP_20_ ligand became bound to half of the FGFR1 cavity, similarly to the binding mode observed for HP_10_ in the crystal structure^66^, while the other HP_20_ ligand interacted partially with the remainder of the cavity as well as FGF2 (Error! Reference source not found.**B**). In the remaining productive replicates, HP_20_ sampled the whole of the binding cavity (**Figure S4**Error! Reference source not found.**A, C**), supporting previous hypotheses for the interaction of longer ligands with the FGFR1-FGF2 complex^71^.

#### 2.8.2 Bikunin-CS proteoglycan

Bikunin PG is a 16 kDa serine protease inhibitor belonging to the kunin family of inhibitors ^76^ and contains a single GAG attached via an O-linked modification at the Ser-10 position^55^. The resultant PG is a therapeutically important molecule that has been used for the treatment of acute pancreatitis^77^. The 3D structure of the GAG sequences in either the free^23,61,78–80^ or bikunin-attached form^61^ have been studied, and found to range from a length of 27 to 39 saccharide units^57^ when attached to bikunin. Therefore, either a 30-mer CS (CS_30_) or a 40-mer CS (CS_40_) chain was used to generate an O-linked bikunin PG via a tetrasaccharide linker (LIN) (**Figure 4**). Subsequent simulations were used to explore the dynamics of these ‘extremes’ of CS chain length, validated against hydrodynamic radii estimated from previously performed DLS measurements.

The R_g_ values were monitored throughout the simulation. We found that starting from an extended conformation, the CS saccharides near the protein start interacting with the protein while the terminal ends point toward the solvent (**Figure 10A, B**). The same was observed while measuring the R_g_ values which were found to gradually reduce throughout the simulation, stabilizing at around ∼3 and ∼4 nm for CS_30_ and CS_40_, respectively. The experimentally estimated R_g_ values (*R*_*g*_ = 1.17 ∗ *R*_ℎ_) lie within the predicted R_g_ values observed for the two simulation extremes, confirming that the most prominent chain length indeed lies within the previously estimated chain length range of 27-39 CS units^57^ (**Figure 10C**).

**Figure 10.**
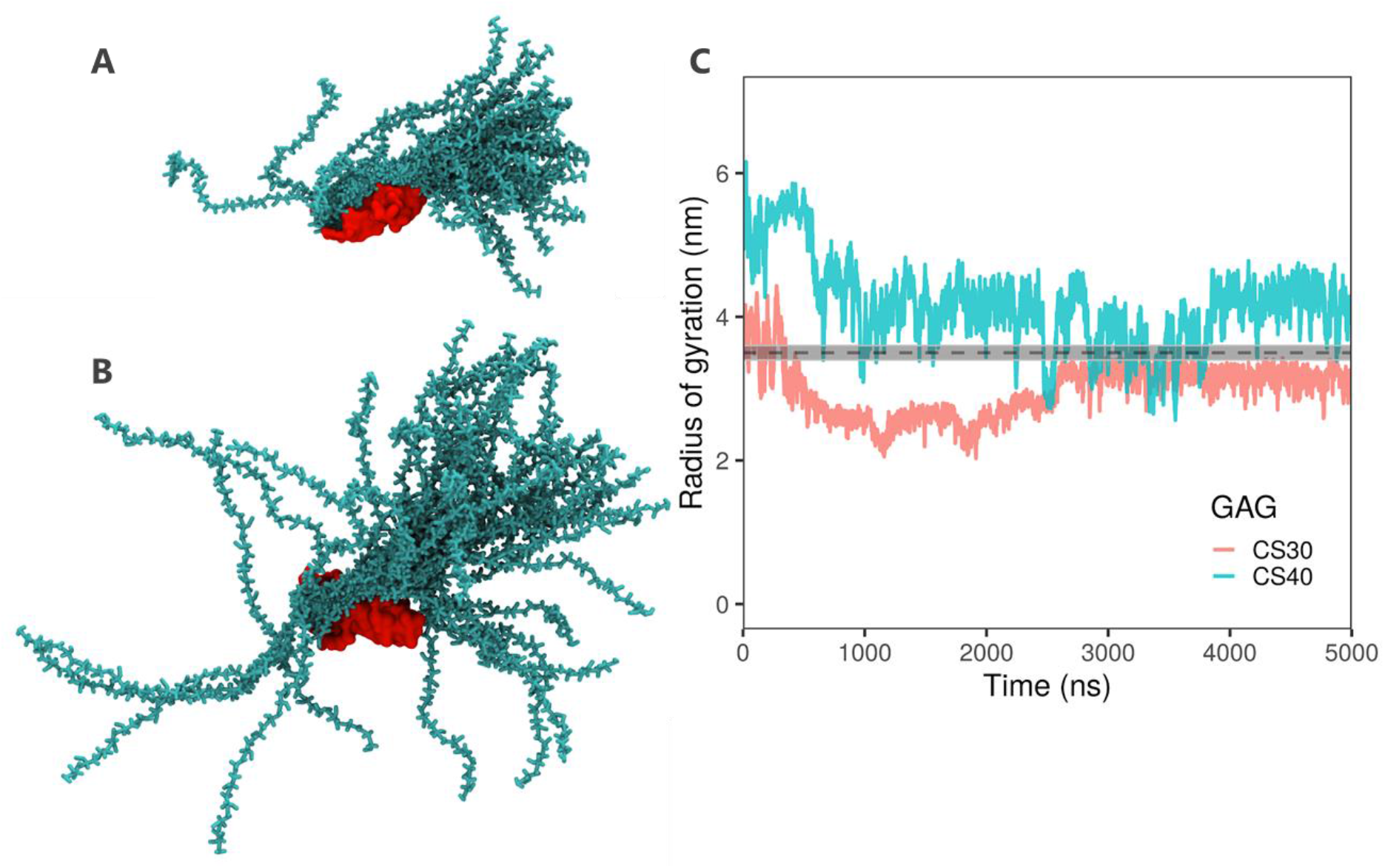
Dynamics of Bikunin PG bound by CS GAG chains. Ensemble of conformations of **(A)** CS_30_ and **(B)** CS_40_ attached to Bikunin. GAG chains are colored in cyan while bikunin is colored in red. Snapshots are overlaid from every 100 ns of the entire trajectory. **(C)** Radius of gyration for bikunin complex formed with CS_30_ (orange) or CS_40_ (cyan). The experimental value of R_g_ is shown as a dashed line with errors in grey. Simulations were performed using Martini v2.2 parameters with a LJ scaling factor of 0.9.

### 2.9 Discussion

Structural characterization of GAGs in PGs is hampered by the fact that their observable lengths in crystal structures are typically short (typically only up to 5 to 8 saccharides)^81^. GAG-protein analyses performed in the past have typically focused on shorter GAG sequences^82–86^. Although these results may be extrapolated to longer polymers, the need for methodologies allowing for directly studying longer sequences has motivated the development of CG GAG models. The CG approach allows for the sampling of extended timescales for larger GAG polymers and PG complexes. Previous CG approaches^23–26^ have made advancements in understanding the behaviour of longer GAG chains, but lack wider compatibility with other classes of biomolecules.

Herein, we report a CG model for a library of common GAG types compatible with the widely popular CG Martini FFs (v2.2 and v3.0) for biomolecular simulations. CG models for HP, HS, CS, DS, KS and HA were carefully developed after assessing the stability of two alternative mapping schemes. A linear mapping scheme similar to previous ones for carbohydrates^30^ and N-glycans^34^ successfully reproduced the local and global properties of tetra- and decasaccharide structures according to AA data for all GAGs, with notable deviations only evident in the case of the flexible KS polymer whose multiple local minima in dihedral space were hard to fully replicate at CG resolution. The conformational properties of the GAGs were also further validated against available experimental hydrodynamic data.

The GAG parameters are still potentially limited by the extent to which anomeric linkages are distinguished in the di/tetrasaccharides. However, this effect is minimized with longer chains where the helical nature is dependent upon the backbone dihedrals and angles. In addition to the glycosidic linkages, the conformations of GAG chains are influenced by the monosaccharide ring conformations, which have been shown to affect local and global properties of glycans, including GAGs^25,87–89^. In our AA simulations (**Figure S5**), the UA and HEX units populated the ^4^*C*_1_ conformation as observed in other FFs as well as NMR studies^87–89^. ^4^*C*_1_ (major) and ^2^*S*_0_ (minor) ring conformers were observed for the GlcA sugar, similar to the lowest energy conformers observed in previously reported MD simulations^88^. The IduA sugar in HP and DS populated the ^1^*C*_4_ conformer, analogous to experimental observations^87^. However, a slight preference towards the ^2^*S*_*o*_ (37%) conformer was seen depending on the sulphonation state of IduA and its neighbouring hexosamine sugars. Hence, the AA FF used reproduces the experimentally observed puckering conformations, increasing the confidence in the obtained CG GAG models. Whilst puckering of individual units cannot be reproduced at the CG resolution, the global puckering dynamics should emerge in the angles and dihedrals along the GAG chains.

The non-bonded parameters governing the interactions of GAGs with water, ions and other biomolecules are equally important^4,90–92^. The Martini bead type choices were validated by comparing partition coefficients to empirical fragment-based prediction methods. Both v2.2 and v3.0 resulted in log *P*_*ow*_ values in the range of the predicted data, meaning that either of the FF versions would be suitable for simulations. On the other hand, in the case of systems containing multiple solvated GAG molecules, we found that the GAGs aggregated within a few hundred nanoseconds for v2.2 of the FF. Similar behaviour has been previously reported wherein the v2.2 FF turned out to be ‘sticky’ especially for carbohydrate containing molecules^33,34,65,93^. Similar to N-glycan^34^ and carbohydrate^65^ studies, a scaling approach proved beneficial for optimizing the nonbonded interactions. Based on the solution aggregate simulations, a scaling factor of 0.9 could be recommended for Martini v2.2, while no scaling should be necessary with Martini v3.0. Following the corrections to the nonbonded interactions, the model performed very well in predictions of protein-GAG interactions and dynamics. Long HS chains were shown to bind spontaneously to FGFR1-FGF2 at the proposed binding site, whilst O-linked CS chains on the bikunin PG could map out the hydrodynamic space for the predicted GAG length. This confirms the model’s applicability to biologically relevant systems.

## 3 Conclusions

In summary, we focused on extending v2.2 and v3.0 of the Martini FF to GAGs. The resultant model can accurately predict local and global properties such as end-to-end distances and hydrodynamic radii. Solvation free energies were in good agreement with prediction data, thereby justifying the choice of bead types. In solution, a modest correction factor of 0.9 was best able to reproduce the underlying atomic resolution *g*(*r*) for Martini v2.2. Applying the parameters to test systems composed of protein-GAG and PG complexes successfully reproduced a range of structural and biophysical data. The model may thus be employed for studying GAG properties in solution or in complex with proteins, benefitting from the efficiency and broad applicability of the Martini CG framework.

## Acknowledgements

The computational work for this article was partially performed on resources of the National Supercomputing Centre, Singapore (https://www.nscc.sg) and BII-ASTAR in house clusters. The research leading to these results was supported by funding from the Ministry of Education in Singapore (MOE AcRF Tier 3 Grant Number MOE2012-T3-1-008) and BII (A*STAR) core funds. PJB acknowledges funding from A*STAR (grant number FY21_CF_HTPO SEED_ID_BII_C211418001). JKM acknowledges AME Young Individual Research Grant (YIRG) number A2084c0160.

