## Supplementary material for "A coarse-grained model of glycosaminoglycans for biomolecular simulations": Supplimenrtary Information

### A coarse-grained model of glycosaminoglycans for biomolecular simulations: Supplementary Information

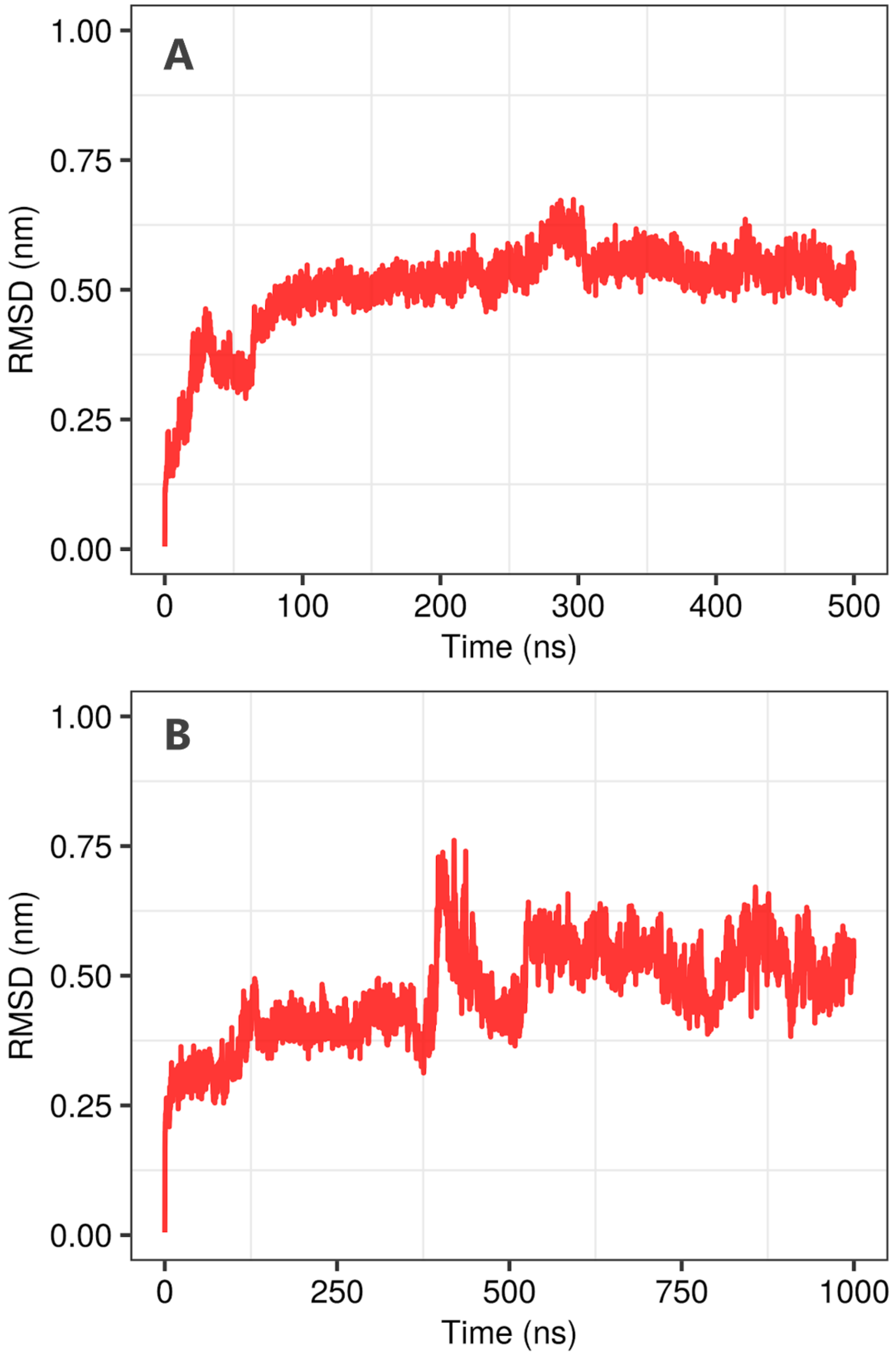

**Figure S1. RMSD during CG simulations of FGFR1-FGF2 and bikunin proteins.** Backbone RMSDs were calculated for **(A)** FGFR1-FGF2 and **(B)** bikunin compared to the respective experimental crystal structures. The higher order protein structure was maintained using an EN with lower and upper cutoffs of 0.5 and 0.9 nm, respectively, with a force constant of 500 kJ $mol^{-1}$ $nm^{-2}$.

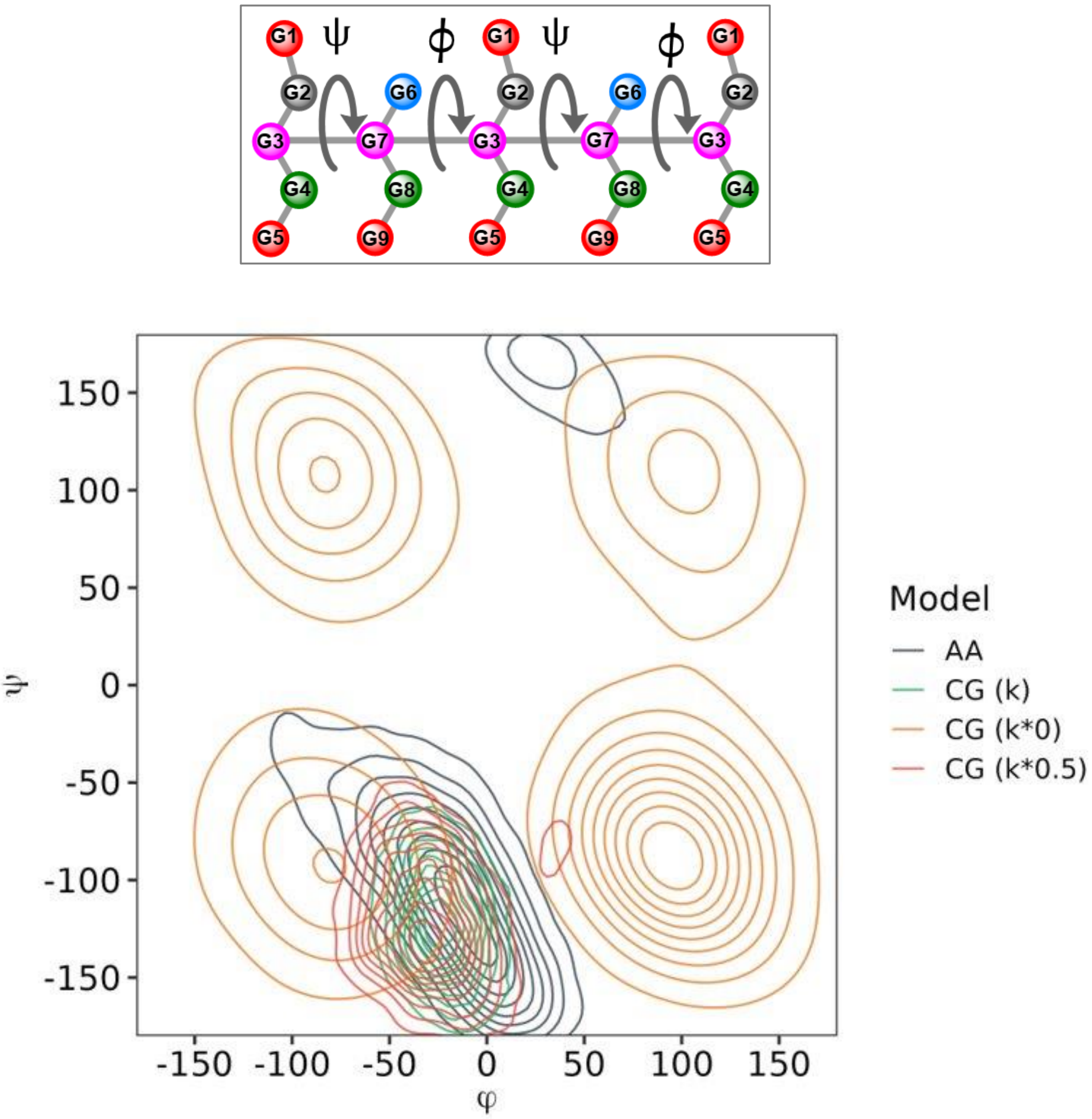

**Figure S2. Effect of backbone dihedral force constant on the flexibility of HP GAG polymer.** The best target CG dihedral potentials (green) resulted in narrower distributions compared to AA data (grey). To test the effect of lowering the dihedral force constants (*k*) upon the distributions, *k*'s of both ϕ and ψ dihedrals were scaled by 50% (i.e. *k**0.5) or switched off (i.e. *k**0). The resultant simulations sample wider distributions compared to unscaled dihedrals but deviate from the global ϕ and ψ minimums.

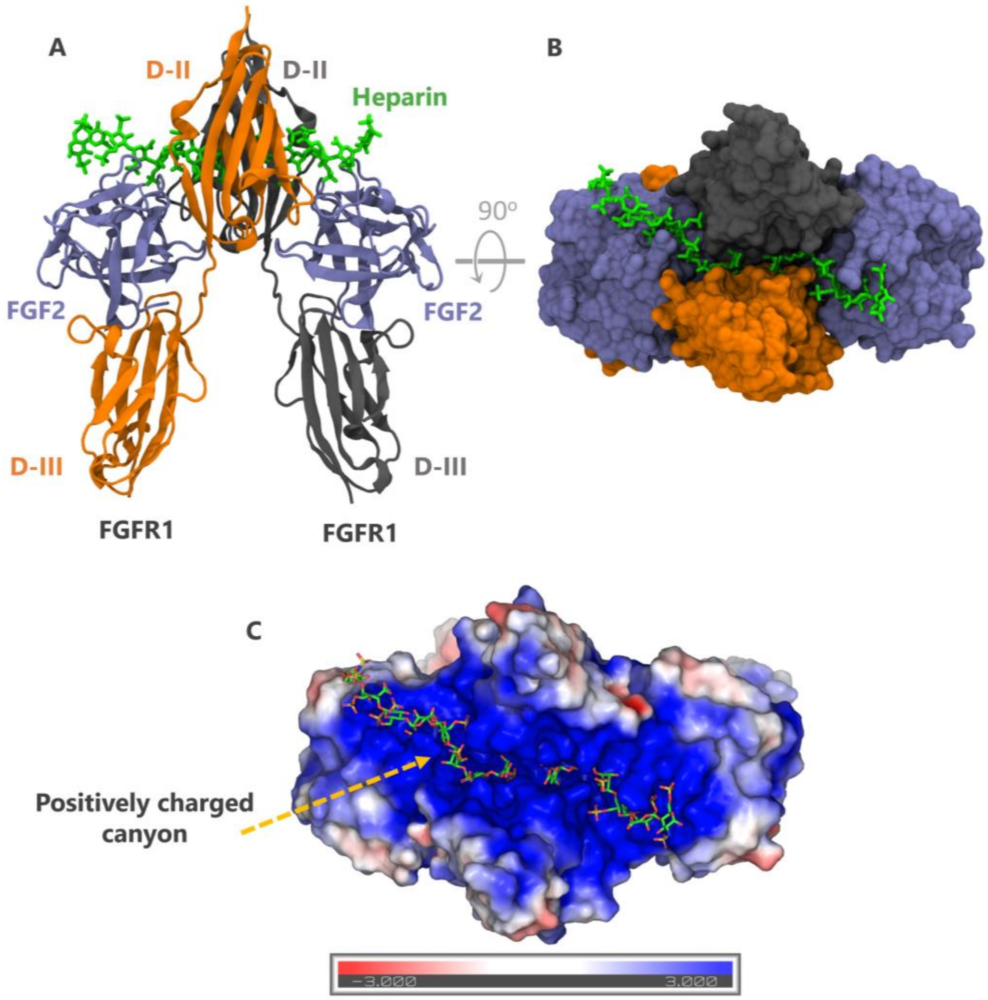

**Figure S3. ECD of FGFR1-FGF2 complex bound by two heparin ($HP_{10}$) molecules, as observed in crystal structure (PDB=1FQ9). (A)** Cartoon representation of complex, with individual protein domains shown in different colors and labelled, and $HP_{10}$ molecules shown in green licorice format. **(B)** Binding interface, with protein shown in surface representation, $HP_{10}$ molecules shown in licorice, and the complex colored as in (A). **(C)** Electrostatic charge distribution of the protein complex (surface representation) highlighting the positively charged canyon which becomes bound by HP (licorice representation). The color-coded electrostatic surface potential of the protein was generated using the Adaptive Poisson-Boltzmann Solver (APBS) with maximum and minimum values set to +3 and -3 $k_BT/e$, respectively.

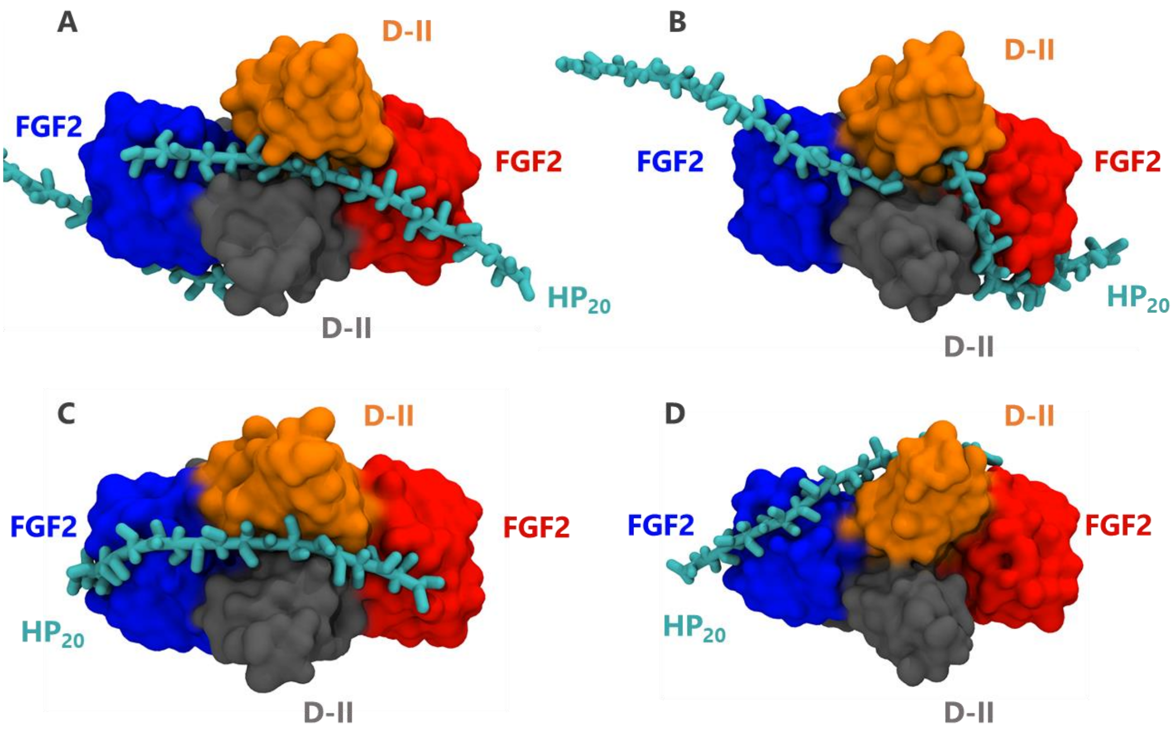

**Figure S4. Unbiased self-assembly simulations of $HP_{20}$ binding to FGFR1-FGF2 complex.** The final simulation snapshot for four replicates is shown in **(A), (B), (C)** and **(D)**. In each figure, the two FGF2 ligands are colored in red and blue while the cavity-forming DII domains on the FGFR1 receptor are colored in orange and grey.

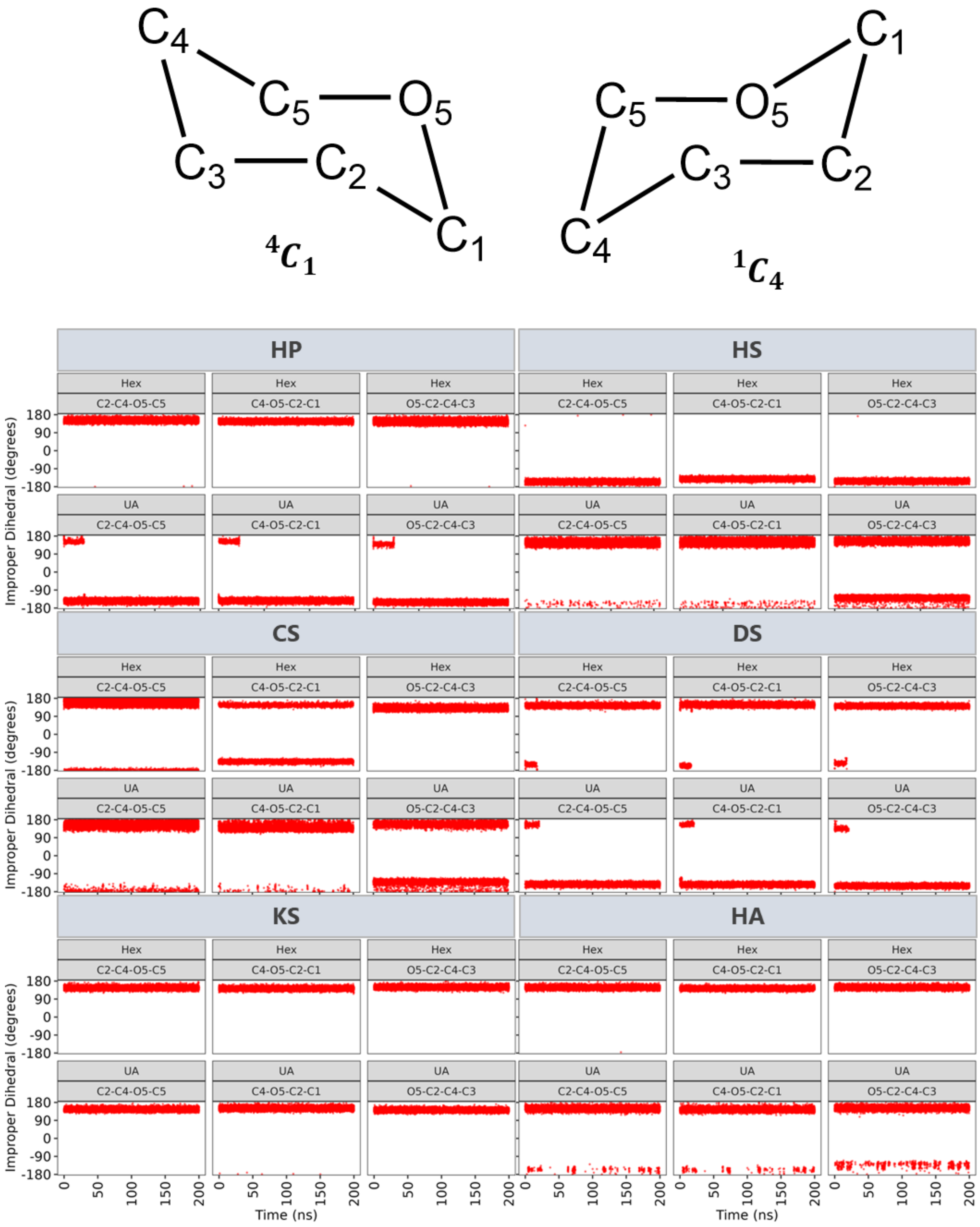

**Figure S5. Dominant ring pucker conformations in the AA GAG models.** Three improper angles C2-C4-O5-C5, C4-O5-C2-C1 and O5-C2-C4-C3 were calculated for each of the Hex and UA sugars of GAG sequences. Two common conformations $^{4}C_{1}$ and $^{1}C_{4}$ are shown. For these conformations, angles are based on the ring pucker nomenclature by Pickett and Strauss[1] and discussed in detail elsewhere[2].

**Table S1. Details of the simulations performed in this study.**

| Parametrization | | |
|---|---|---|
| System | AA | CG |
| Tetrasaccharides (AA) | 500 ns x 6 | 500 ns x 6 |
| Decasaccharides (AA) | 300 ns x 6 | 500 ns x 6 |
| Thermodynamic integration calculations | | |
| System | AA | CG |
| Disaccharides in octanol | 5 ns x ≥ 55 windows | 40 ns x ≥ 55 windows |
| Disaccharides in water | 5 ns x ≥ 55 windows | 40 ns x ≥ 55 windows |
| Protein simulations | | |
| System | CG | |
| FGFR1-FGF2 dimer + $HP_{20}$ (Docked) | 500 ns | |
| FGFR1-FGF2 dimer + $HP_{20}$ (Self-assembly) | 4 x 500 ns | |
| Bikunin + $CS_{30}$ | 5,000 ns | |
| Bikunin + $CS_{40}$ | 5,000 ns | |

**Table S2. Bonded parameters for the various GAG sequences.** Beads are named according to Figure 4 in the main article. Bead names such as G3' represent those from the neighboring disaccharide.

| **Heparin (HP)** | | | | |
|---|---|---|---|---|
| Bonds: atom1; atom2; $r_{bond}$ (nm); $k_{bond}$ (kJ $mol^{-1}nm^{-2}$) | | | | |
| G1 | G2 | 0.32 | 25,000 | |
| G2 | G3 | 0.22 | 25,000 | |
| G3 | G4 | 0.22 | 25,000 | |
| G4 | G5 | 0.31 | 25,000 | |
| G3 | G7 | 0.5 | 25,000 | |
| G6 | G7 | 0.22 | 25,000 | |
| G7 | G8 | 0.22 | 25,000 | |
| G8 | G9 | 0.32 | 15,000 | |
| G7 | G3' | 0.49 | 25,000 | |
| Angles: atom1; atom2; atom3; $\theta_{angle}$ (deg), $k_{angle}$ (kJ $mol^{-1}rad^{-2}$) | | | | |
| G1 | G2 | G3 | 150 | 180 |
| G2 | G3 | G4 | 110 | 150 |
| G3 | G4 | G5 | 100 | 210 |
| G2 | G3 | G7 | 110 | 150 |
| G4 | G3 | G7 | 115 | 250 |
| G6 | G7 | G8 | 125 | 400 |
| G7 | G8 | G9 | 100 | 150 |
| G6 | G7 | G3' | 105 | 170 |
| G8 | G7 | G3' | 120 | 180 |
| G3 | G7 | G3' | 175 | 140 |

| | | | | | | |
|---|---|---|---|---|---|---|
| G7 | G3' | G7' | 175 | 500 | | |
| Dihedrals: atom1; atom2; atom3; atom4; $\theta_{dihedral}$ (deg), $k_{dihedral}$ (kJ $\mathrm{mol}^{-1}$), multiplicity | | | | | | |
| G1 | G2 | G3 | G4 | 110 | 15 | 1 |
| G2 | G3 | G4 | G5 | 180 | 10 | 1 |
| G2 | G3 | G7 | G6 | 100 | 20 | 1 |
| G6 | G7 | G8 | G9 | 40 | 35 | 1 |
| G6 | G7 | G3' | G2' | 90 | 35 | 1 |
| G3 | G7 | G3' | G7' | 130 | 5 | 1 |
| **Heparan Sulphate (HS)** | | | | | | |
| Bonds: atom1; atom2; $r_{bond}$ (nm); $k_{bond}$ (kJ $\mathrm{mol}^{-1}\mathrm{nm}^{-2}$) | | | | | | |
| G1 | G2 | 0.32 | 30,000 | | | |
| G2 | G3 | 0.21 | 25,000 | | | |
| G3 | G4 | 0.21 | 25,000 | | | |
| G4 | G5 | 0.34 | 25,000 | | | |
| G4 | G6 | 0.245 | 25,000 | | | |
| G7 | G8 | 0.21 | 25,000 | | | |
| G8 | G9 | 0.21 | 30,000 | | | |
| G9 | G10 | 0.3 | 8,000 | | | |
| G3 | G8 | 0.525 | 25,000 | | | |
| G8 | G3' | 0.475 | 10,000 | | | |
| Angles: atom1; atom2; atom3; $\theta_{angle}$ (deg), $k_{angle}$ (kJ $\mathrm{mol}^{-1}\mathrm{rad}^{-2}$) | | | | | | |
| G1 | G2 | G3 | 135 | 160 | | |
| G2 | G3 | G4 | 120 | 280 | | |
| G3 | G4 | G5 | 148 | 200 | | |
| G3 | G4 | G6 | 70 | 170 | | |
| G5 | G4 | G6 | 135 | 120 | | |
| G7 | G8 | G9 | 135 | 280 | | |
| G8 | G9 | G10 | 160 | 100 | | |
| G2 | G3 | G8 | 100 | 260 | | |
| G4 | G3 | G8 | 110 | 240 | | |
| G7 | G8 | G3' | 90 | 50 | | |
| G9 | G8 | G3' | 140 | 250 | | |
| G3 | G8 | G3' | 175 | 500 | | |
| G8 | G3' | G8' | 160 | 150 | | |
| Dihedrals: atom1; atom2; atom3; atom4; $\theta_{dihedral}$ (deg), $k_{dihedral}$ (kJ $\mathrm{mol}^{-1}$, multiplicity | | | | | | |
| G1 | G2 | G3 | G4 | 30 | 25 | 1 |
| G2 | G3 | G4 | G5 | 200 | 25 | 1 |
| G2 | G3 | G8 | G7 | 70 | 25 | 1 |
| G7 | G8 | G3' | G2' | 120 | 20 | 1 |
| G3 | G8 | G3' | G8' | 280 | 2 | 1 |
| G8 | G3' | G8' | G3" | 310 | 7 | 1 |
| **Chondroitin Sulphate (CS)** | | | | | | |
| Bonds: atom1; atom2; $r_{bond}$ (nm); $k_{bond}$ (kJ $\mathrm{mol}^{-1}\mathrm{nm}^{-2}$) | | | | | | |

| | | | | | | |
|---|---|---|---|---|---|---|
| G1 | G2 | 0.325 | 25,000 | | | |
| G2 | G3 | 0.2 | 25,000 | | | |
| G3 | G4 | 0.2 | 30,000 | | | |
| G4 | G5 | 0.325 | 2,000 | | | |
| G3 | G6 | 0.295 | 30,000 | | | |
| G3 | G8 | 0.595 | 10,000 | | | |
| G7 | G8 | 0.22 | 25,000 | | | |
| G8 | G9 | 0.2 | 25,000 | | | |
| G9 | G10 | 0.295 | 15,000 | | | |
| G8 | G3' | 0.25 | 5,000 | | | |
| Angles: atom1; atom2; atom3; $\theta_{angle}$ (deg), $k_{angle}$ (kJ $mol^{-1}rad^{-2}$) | | | | | | |
| G1 | G2 | G3 | 132 | 200 | | |
| G2 | G3 | G4 | 95 | 150 | | |
| G3 | G4 | G5 | 130 | 200 | | |
| G2 | G3 | G6 | 80 | 170 | | |
| G4 | G3 | G6 | 120 | 200 | | |
| G7 | G8 | G9 | 80 | 100 | | |
| G8 | G9 | G10 | 105 | 80 | | |
| G8 | G3' | G2' | 50 | 80 | | |
| G8 | G3' | G4' | 110 | 350 | | |
| G3 | G8 | G7 | 70 | 150 | | |
| G3 | G8 | G9 | 75 | 230 | | |
| Dihedrals: atom1; atom2; atom3; atom4; $\theta_{dihedral}$ (deg), $k_{dihedral}$ (kJ $mol^{-1}$), multiplicity | | | | | | |
| G1 | G2 | G3 | G4 | -20 | 25 | 1 |
| G2 | G3 | G4 | G5 | 280 | 30 | 1 |
| G7 | G8 | G9 | G10 | 10 | 30 | 1 |
| G2 | G3 | G8 | G7 | 140 | 30 | 1 |
| G7 | G8 | G3' | G2' | -30 | 30 | 1 |
| **Dermatan Sulphate (DS)** | | | | | | |
| Bonds: atom1; atom2; $r_{bond}$ (nm); $k_{bond}$ (kJ $mol^{-1}nm^{-2}$) | | | | | | |
| G1 | G2 | 0.222 | 20,000 | | | |
| G2 | G3 | 0.222 | 20,000 | | | |
| G3 | G4 | 0.297 | 22,000 | | | |
| G2 | G5 | 0.29 | 30,000 | | | |
| G2 | G7 | 0.57 | 20,000 | | | |
| G6 | G7 | 0.22 | 30,000 | | | |
| G7 | G8 | 0.2 | 20,000 | | | |
| G8 | G9 | 0.315 | 30,000 | | | |
| G7 | G2' | 0.48 | 35,000 | | | |
| Angles: atom1; atom2; atom3; $\theta_{angle}$ (deg), $k_{angle}$ (kJ $mol^{-1}rad^{-2}$) | | | | | | |
| G1 | G2 | G3 | 143 | 200 | | |
| G2 | G3 | G4 | 136 | 250 | | |
| G1 | G2 | G5 | 80 | 180 | | |
| G1 | G2 | G7 | 120 | 230 | | |

| | | | | | | |
|---|---|---|---|---|---|---|
| G3 | G2 | G7 | 30 | 250 | | |
| G6 | G7 | G8 | 130 | 450 | | |
| G7 | G8 | G9 | 102 | 270 | | |
| G6 | G7 | G2' | 113 | 300 | | |
| G8 | G7 | G2' | 112 | 370 | | |
| G2 | G7 | G2' | 165 | 330 | | |
| G7 | G2' | G7' | 115 | 450 | | |
| Dihedrals: atom1; atom2; atom3; atom4; $\theta_{dihedral}$ (deg), $k_{dihedral}$ (kJ $mol^{-1}$, multiplicity | | | | | | |
| G1 | G2 | G3 | G4 | 280 | 30 | 1 |
| G1 | G2 | G7 | G6 | 205 | 35 | 1 |
| G6 | G7 | G8 | G9 | 45 | 35 | 1 |
| G2 | G7 | G2' | G7' | 70 | 40 | 1 |
| G7 | G2' | G7' | G2" | 200 | 30 | 1 |
| **Keratan Sulphate (KS)** | | | | | | |
| Bonds: atom1; atom2; $r_{bond}$ (nm); $k_{bond}$ (kJ $mol^{-1}nm^{-2}$) | | | | | | |
| G1 | G2 | 0.222 | 30,000 | | | |
| G2 | G3 | 0.222 | 30,000 | | | |
| G3 | G4 | 0.222 | 30,000 | | | |
| G4 | G5 | 0.312 | 30,000 | | | |
| G6 | G7 | 0.325 | 30,000 | | | |
| G7 | G8 | 0.222 | 30,000 | | | |
| G8 | G9 | 0.222 | 30,000 | | | |
| G3 | G8 | 0.495 | 30,000 | | | |
| G8 | G3' | 0.605 | 30,000 | | | |
| Angles: atom1; atom2; atom3; $\theta_{angle}$ (deg), $k_{angle}$ (kJ $mol^{-1}rad^{-2}$) | | | | | | |
| G1 | G2 | G3 | 120 | 200 | | |
| G2 | G3 | G4 | 125 | 350 | | |
| G3 | G4 | G5 | 150 | 200 | | |
| G6 | G7 | G8 | 135 | 230 | | |
| G7 | G8 | G9 | 130 | 280 | | |
| G2 | G3 | G8 | 120 | 300 | | |
| G4 | G3 | G8 | 105 | 380 | | |
| G7 | G8 | G3' | 125 | 280 | | |
| G9 | G8 | G3' | 35 | 200 | | |
| G3 | G8 | G3' | 115 | 200 | | |
| G8 | G3' | G8' | 170 | 350 | | |
| Dihedrals: atom1; atom2; atom3; atom4; $\theta_{dihedral}$ (deg), $k_{dihedral}$ (kJ $mol^{-1}$, multiplicity | | | | | | |
| G1 | G2 | G3 | G4 | 45 | 20 | 1 |
| G2 | G3 | G4 | G5 | 170 | 20 | 1 |
| G6 | G7 | G8 | G9 | 280 | 5 | 1 |
| G2 | G3 | G8 | G7 | 70 | 20 | 1 |
| G7 | G8 | G3' | G2' | 160 | 30 | 1 |
| G3 | G8 | G3' | G8' | 210 | 6 | 1 |

| | | | | | | |
|---|---|---|---|---|---|---|
| **Hyaluronic acid (HA)** | | | | | | |
| Bonds: atom1; atom2; $r_{bond}$ (nm); $k_{bond}$ (kJ $mol^{-1}nm^{-2}$) | | | | | | |
| G1 | G2 | 0.222 | 30,000 | | | |
| G2 | G3 | 0.222 | 30,000 | | | |
| G3 | G4 | 0.3 | 30,000 | | | |
| G5 | G6 | 0.222 | 30,000 | | | |
| G6 | G7 | 0.222 | 30,000 | | | |
| G2 | G6 | 0.55 | 25,000 | | | |
| G6 | G2' | 0.525 | 30,000 | | | |
| Angles: atom1; atom2; atom3; $\theta_{angle}$ (deg), $k_{angle}$ (kJ $mol^{-1}rad^{-2}$) | | | | | | |
| G1 | G2 | G3 | 120 | 400 | | |
| G2 | G3 | G4 | 160 | 250 | | |
| G1 | G2 | G6 | 165 | 170 | | |
| G3 | G2 | G6 | 55 | 180 | | |
| G5 | G6 | G7 | 125 | 220 | | |
| G5 | G6 | G2' | 105 | 400 | | |
| G7 | G6 | G2' | 110 | 400 | | |
| G2 | G6 | G2' | 155 | 220 | | |
| G6 | G2' | G6' | 130 | 320 | | |
| Dihedrals: atom1; atom2; atom3; atom4; $\theta_{dihedral}$ (deg), $k_{dihedral}$ (kJ $mol^{-1}$), multiplicity | | | | | | |
| G1 | G2 | G3 | G4 | 195 | 20 | 1 |
| G3 | G2 | G6 | G7 | 220 | 50 | 1 |
| G7 | G6 | G2' | G3' | 30 | 50 | 1 |
| G2 | G6 | G2' | G6' | 260 | 20 | 1 |
| G6 | G2' | G6' | G2' | 0 | 10 | 1 |
| **GAG Linker (LIN)** | | | | | | |
| Bonds: atom1; atom2; $r_{bond}$ (nm); $k_{bond}$ (kJ $mol^{-1}nm^{-2}$) | | | | | | |
| SC | G2 | 0.43 | 25,000 | | | |
| G1 | G2 | 0.22 | 25,000 | | | |
| G2 | G3 | 0.22 | 20,000 | | | |
| G2 | G5 | 0.475 | 15,000 | | | |
| G4 | G5 | 0.22 | 20,000 | | | |
| G5 | G6 | 0.22 | 20,000 | | | |
| G5 | G8 | 0.54 | 10,000 | | | |
| G7 | G8 | 0.22 | 20,000 | | | |
| G8 | G9 | 0.22 | 25,000 | | | |
| G8 | G11 | 0.6 | 15,000 | | | |
| G10 | G11 | 0.22 | 25,000 | | | |
| G11 | G12 | 0.22 | 25,000 | | | |
| Angles: atom1; atom2; atom3; $\theta_{angle}$ (deg), $k_{angle}$ (kJ $mol^{-1}rad^{-2}$) | | | | | | |
| BB | SC | G2 | 135 | 140 | | |
| G0 | G2 | G1 | 20 | 300 | | |
| G0 | G2 | G3 | 30 | 300 | | |
| G0 | G2 | G5 | 175 | 150 | | |

| | | | | | | |
|---|---|---|---|---|---|---|
| G2 | G5 | G8 | 150 | 170 | | |
| G5 | G8 | G11 | 140 | 150 | | |
| G1 | G2 | G3 | 75 | 500 | | |
| G4 | G5 | G6 | 130 | 280 | | |
| G7 | G8 | G9 | 130 | 320 | | |
| G10 | G11 | G12 | 130 | 370 | | |
| G1 | G2 | G5 | 140 | 100 | | |
| G3 | G2 | G5 | 170 | 200 | | |
| G4 | G5 | G8 | 140 | 130 | | |
| G6 | G5 | G8 | 40 | 150 | | |
| G7 | G8 | G11 | 125 | 150 | | |
| G9 | G8 | G11 | 60 | 150 | | |
| Dihedrals: atom1; atom2; atom3; atom4; $\theta_{dihedral}$ (deg), $k_{dihedral}$ (kJ mol$^{-1}$), multiplicity | | | | | | |
| BB | SC | G2 | G5 | 170 | 10 | 1 |
| G0 | G2 | G5 | G8 | 250 | 30 | 1 |
| G2 | G5 | G8 | G11 | 220 | 10 | 1 |
| G1 | G2 | G5 | G4 | 80 | 30 | 1 |
| G4 | G5 | G8 | G7 | 200 | 20 | 1 |
| G7 | G8 | G11 | G10 | 160 | 15 | 1 |
| G-1 | G0 | G2 | G3 | 40 | 25 | 1 |

**Table S3. Comparison of partition coefficients obtained from CG simulations and various empirical prediction methods.** Both Martini v2.2 and v3.0 FFs were used for the calculations. The values from different empirical prediction methods were averaged to compare with the simulation results.

| GAG | Martini FF | $P_{ow}$ (Sim) | Predicted (Average) | ClogP | AC_logP | ALOGP | XLOGP3 |
|---|---|---|---|---|---|---|---|
| HP | v3.0 | -8.5 | -8.1 | -9.7 | -10.8 | -6.1 | -6.0 |
| | v2.2 | -6.9 | | | | | |
| HS | v3.0 | -9.1 | -8.8 | -9.3 | -12.4 | -7.0 | -6.7 |
| | v2.2 | -7.5 | | | | | |
| CS | v3.0 | -7.6 | -5.7 | -6.4 | -6.6 | -4.9 | -4.9 |
| | v2.2 | -4.0 | | | | | |
| DS | v3.0 | -6.6 | -5.7 | -6.4 | -6.6 | -4.9 | -4.9 |
| | v2.2 | -4.0 | | | | | |
| KS | v3.0 | -7.3 | -6.9 | -7.7 | -8.2 | -5.7 | -6.1 |
| | v2.2 | -6.0 | | | | | |
| HA | v3.0 | -5.0 | -4.4 | -4.8 | -4.6 | -4.3 | -3.9 |
| | V2.2 | -4.4 | | | | | |

**Movie S1: Effect of dihedrals from tetrasaccharides vs decasaccharides.** The movie shows the helical GAG chains upon incorporation of backbone dihedrals from decasaccharide all atom simulations.
